# Circadian modulation of mosquito host-seeking persistence by Pigment-Dispersing Factor impacts daily biting patterns

**DOI:** 10.1101/2024.09.19.613886

**Authors:** Linhan Dong, Richard Hormigo, Jord M. Barnett, Chloe Greppi, Laura B. Duvall

## Abstract

Daily rhythms in mosquito attraction to humans are thought to drive biting patterns and contribute to disease transmission dynamics. Behavioral rhythms in many insects are controlled by specialized clock cells in the brain that are coordinated by the neuropeptide Pigment-Dispersing Factor (PDF). We show that female *Aedes aegypti* mosquitoes with genetically disrupted PDF display altered daily behavioral timing with reduced locomotor activity and biting in the morning. Using an automated behavioral tracking system, we also report that mosquitoes exhibit daily modulation of response persistence to the host cue carbon dioxide and loss of PDF alters this pattern. These findings indicate that PDF regulates temporal features of host-seeking behavior that promote biting success at specific times of day and may underlie blood feeding patterns observed in the field.

---

Female mosquitoes bite and consume a blood meal to obtain nutrition for egg development. When blood feeding from human hosts, they can transmit pathogens that cause diseases including malaria, dengue, Zika, and West Nile fever (*1*–*3*). However, biting does not occur uniformly throughout the day, species-specific rhythms maximize biting success while avoiding unfavorable environmental conditions and predators in their ecological niche (*4*–*8*). These daily rhythms play critical roles in disease transmission dynamics and understanding the temporal patterns of mosquito/host interactions is important for effective application of vector control approaches to target mosquitoes during periods of active biting (*9, 10*).

Although daily rhythms in behavior and physiology can entrain to environmental cues like dawn and dusk, they are sustained in the absence of environmental cues by endogenous pacemakers (*11*–*13*). These rhythms are controlled by the regulated expression of specialized circadian genes and proteins in critical pacemaker cells in the brain. These “clock” cells express autoregulatory transcription and translation feedback loops (TTFL) that coordinate physiological and behavioral rhythms (*14*). The clock genes that constitute the TTFL are well characterized in *Drosophila* and largely conserved in mosquitoes (*15*–*17*). Recent studies have shown that whole-body knock-outs or RNAi-mediated knockdown of core components of the mosquito TTFL lead to loss of daily behavioral patterns and circadian rhythmicity (*18*–*20*).

In insects, dozens to hundreds of individual clock cells in the brain form anatomically and functionally distinct subgroups (*21*) that must be coordinated for the expression of coherent organismal rhythms (*22, 23*). Intercellular coordination is achieved through the action of Pigment-Dispersing Factor (PDF), a neuropeptide produced in some clock cell subgroups that acts as a critical synchronizing agent to organize molecular, cellular, and behavioral rhythms (*24*–*27*). In *Aedes aegypti*, PDF is produced by groups of small and large ventral cells that co-express the core clock protein PERIOD (*28*).

Some rhythms can function independently of the central brain by relying on clock gene expression in peripheral tissues. In *Drosophila*, olfactory responses are regulated by antennal clocks that require intact TTFL genes but do not require PDF (*29, 30*). Mosquito host-seeking requires coordinated locomotor responses to host-related chemosensory cues including carbon dioxide, human odor, and body temperature (*31*–*41*). Carbon dioxide (CO_2_) is a critical host-seeking cue that functions as a behavioral activator that triggers a persistent predatory state that heightens the saliency of other host cues, allowing for the integration of multimodal sensory signals, ultimately leading to successful biting (*32, 34, 40, 41*). Although it is clear that chemosensory processing is required for mosquito host-seeking and biting and previous work has shown that olfactory sensitivity is modulated throughout the day (*42*), we lack a mechanistic view as to how behavioral responses to host cues vary at different times of day and whether these responses depend on PDF coordination of central brain clock cells. Here, we use videography and flight tracking to characterize time-of-day dependent features of mosquito locomotor activity and CO_2_ responsiveness and we generated PDF-null mutants to explore how central clock disruption affects mosquito activity, host-seeking, and biting.

## PDF regulates daily spontaneous locomotor activity in *Aedes aegypti* females

To determine the role of PDF in controlling daily locomotor activity, we used a genetic approach to disrupt the *pdf* locus and provide genetic access to label cells that normally express the *pdf* gene using the Q-binary system (*43*). To generate a “knock-in/knock-out” line, we used CRISPR-Cas9 tools to insert a cassette that includes the QF2 transcriptional activator (*43, 44*) as well as an eye-specific fluorescent reporter directly into the coding sequence of the *pdf* gene (**fig. S1A–C**). This leads to an early truncation of the *pdf* locus prior to the start of the mature peptide and a complete loss of PDF immunostaining, indicating that this is a null allele (**fig. S1D**). The integration of QF2 also enables *pdf*-specific labeling when crossed with a QUAS-driven fluorescent reporter (*45*) (**fig. S1E**).

To ask how loss of PDF affects locomotor activity, we used video-tracking of individual animals to quantify spontaneous locomotor activity under 12:12 light/dark (LD) conditions with a 30-minute gradual dawn and dusk providing zeitgeber cues (*46*) (**Fig. 1A**). Consistent with previous reports (*42, 46*–*48*), we find that wildtype and *pdf* heterozygote females show a bimodal pattern with morning and evening peaks of activity and a midday siesta period of low activity under LD conditions (**Fig. 1B and C**). As has been reported (*42, 46*), *Aedes aegypti* females show no anticipatory activity before dawn. In homozygous *pdf* mutant females, we observe a dramatic loss of morning activity (**Fig. 1B–D**). The residual morning activity in *pdf* homozygotes occurs with a delayed phase compared to the control groups (**Fig. 1E**). Although the evening peak of activity is maintained in *pdf* mutants, we observe an early termination of activity after dusk, with mutants becoming inactive within 3.23 ± 0.50 minutes of lights-off, compared to wildtype and heterozygous animals (20.60 ± 2.37 minutes and 13.25 ± 1.04 minutes, respectively) (**fig. S2A–B**). These findings indicate that *pdf* is a critical regulator of daily spontaneous locomotor activity under LD conditions and that PDF normally promotes morning locomotor activity, advances the morning peak, and maintains activity for ∼20 minutes after lights-off.

**Figure 1.**
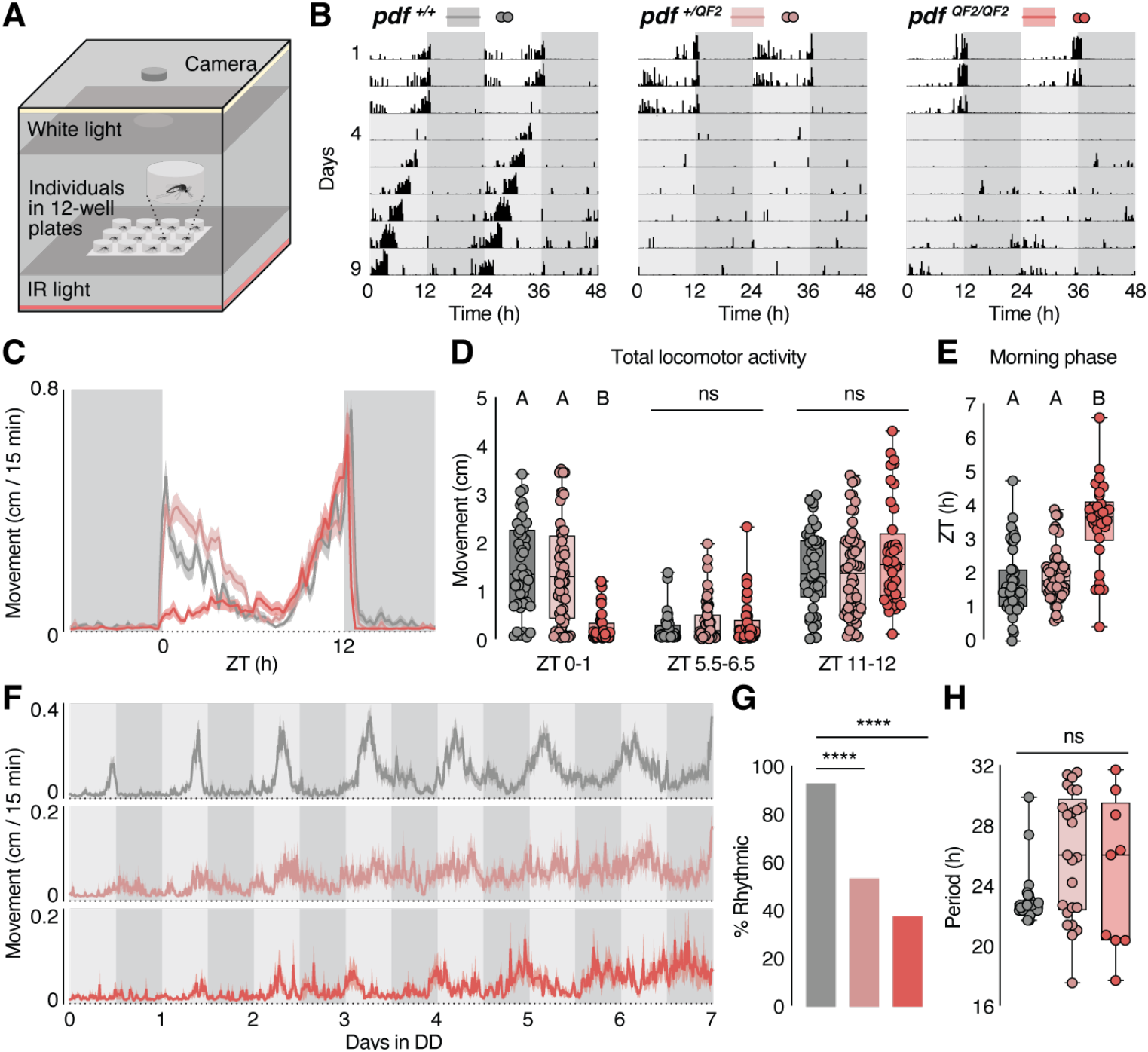
*Pdf* mutants show deficits in morning locomotor activity and circadian rhythmicity. (**A**) Schematic of the MozzieBox locomotor activity assay. Female mosquitoes were loaded individually into the wells of 12-well plates, and infrared images were taken every minute to track mosquito movement. (**B**) Representative double-plotted actograms of female mosquitoes in the locomotor activity assay. Activity shown in 15-minute bins. Locomotor activity was recorded under three days of 12 hours light: 12 hours dark LD cycles with 30 min dawn/dusk light transition periods (white and dark grey shading represent light and dark phases, respectively), followed by seven days of DD cycles (light grey shading represents respective light phase from prior LD entrainment). (**C**) Activity profiles under LD entrainment. Activities from ZT18 of the second LD cycle to ZT18 of the third LD cycle are shown. For all analysis associated with LD entrainment (C–E), n = 43 for *pdf* ^*+/+*^, n = 53 for *pdf* ^*+/QF2*^, and n = 40 for *pdf* ^*QF2/QF2*^. (**D**) Total locomotor activity during ZT0–1, ZT5.5–6.5, and ZT11–12. Kruskal-Wallis test with Dunn’s multiple comparisons. Columns labeled with different letters are significantly different from each other (P < 0.0001). (**E**) Phase of morning locomotor activity. Phase was quantified after applying a Savitzky-Golay filter (see Methods). Ordinary one-way ANOVA with Tukey’s multiple comparisons test. Columns labeled with different letters are significantly different from each other (P < 0.0001). (**F**) Activity profiles in DD. For analysis associated with DD (F–G), n = 40 for *pdf* ^*+/+*^, n = 47 for *pdf* ^*+/QF2*^, and n = 24 for *pdf* ^*QF2/QF2*^. (**G**) Percentage of rhythmic animals under DD. Animal were considered rhythmic if rhythmic power > 50 (Chi-square periodogram) Chi-square test. (**H**) Chi-square periodogram analysis of free-running period (n = 37 for *pdf* ^*+/+*^, n = 25 for *pdf* ^*+/QF2*^, and n = 9 for *pdf* ^*QF2/QF2*^). Arrhythmic animals were excluded from period length analysis. Kruskal-Wallis test with Dunn’s multiple comparisons. ****P < 0.0001; P > 0.05: not significant (ns). Shaded lines indicate mean ± SEM. Dots represent individual animals. Boxes extend from the 25th to 75th percentiles.

## PDF regulates circadian locomotor rhythmicity

One hallmark of circadian rhythms is that they are endogenously maintained without rhythmic environmental inputs (*11*). Wildtype *Aedes aegypti* females maintain rhythmic activity under constant dark (DD) conditions for >7 days with a period of 22.61 ± 0.08 hours, although the morning peak is lost and activity becomes unimodal (**Fig. 1F–H**). In general, all animals show low levels of locomotor activity during the first day in DD with activity levels gradually increasing over time (**Fig. 1F**). Interestingly, both heterozygous and homozygous *pdf* mutants show high levels of arrhythmicity under constant dark conditions (**Fig. 1G**). Overall, the heterozygous animals show higher levels of phase synchrony during the first days in DD compared to the homozygous mutants, suggesting that the heterozygotes show an intermediate phenotype (**fig. S2C**). These findings indicate that PDF is a crucial regulator of circadian locomotor rhythmicity and behavioral effects associated with the loss of a single copy of the *pdf* gene become apparent under constant dark conditions, in the absence of environmental cues to entrain rhythms.

## Loss of PDF leads to reduced biting efficiency in the morning

Under LD conditions, *pdf* mutants showed a significant reduction in morning activity, while evening activity was less affected. To determine how spontaneous locomotor timing relates to host-cue-elicited biting, we assessed blood feeding at morning and evening timepoints using a standardized combination of host-associated cues. Female mosquitoes were placed in an environmentally-controlled cabinet and exposed to a 30 second pulse of 10% CO_2_, which was allowed to dissipate for 1.5 minutes to increase levels of CO_2_ by ∼1,000 ppm and the cabinet door was then opened to allow CO_2_ levels to return to baseline before offering sheep blood in an artificial feeder baited with human host odor and heated to 37°C. Females were allowed 10 minutes to feed and then scored for the presence of fresh blood in their abdomen (**Fig. 2A**). Wildtype and heterozygous females reliably blood feed in the morning (ZT0–1) and evening (ZT11–12) with highest levels of feeding in the evening (ZT11–12). Homozygous *pdf* mutant females showed no significant differences in evening feeding compared to other genotypes, although the mutants showed higher variation at this timepoint, but blood feeding was significantly reduced in the morning (**Fig. 2B**), at the same time when *pdf* mutants also show significantly reduced spontaneous locomotor activity. This suggests that behavioral timing changes induced by loss of PDF also affect biting efficiency in a time-of-day specific manner.

**Figure 2.**
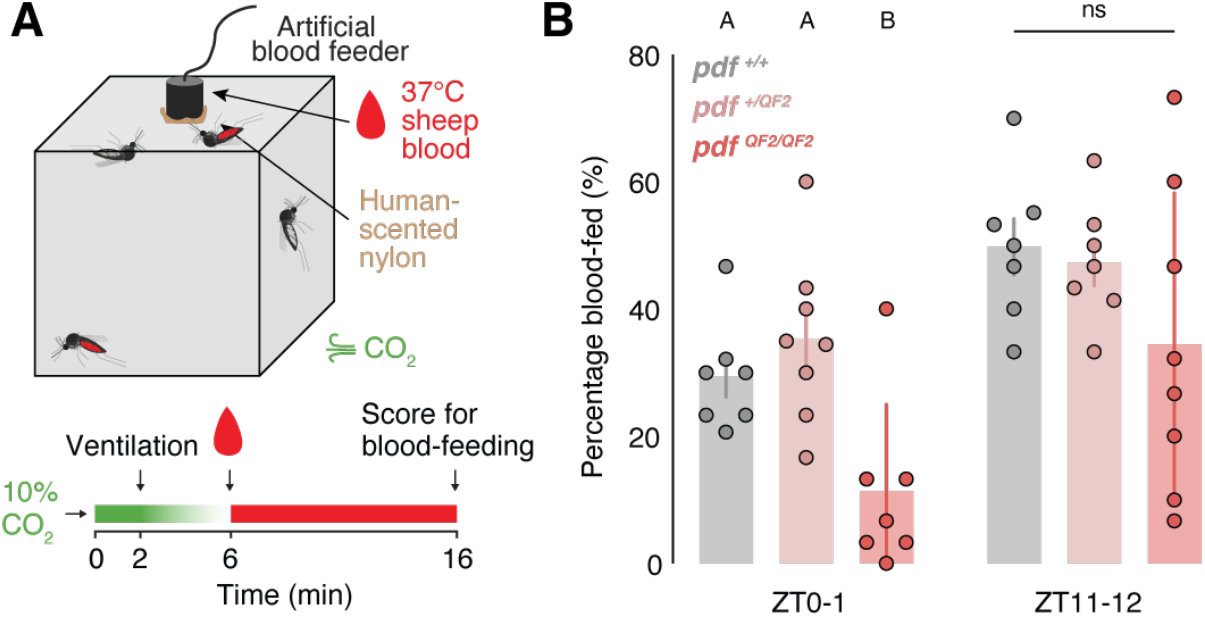
*Pdf* mutants show reduced blood feeding in the morning. (**A**) Schematic of the blood-feeding assay. Mosquitoes in a 24.5 cm cubical cage were provided with host-associated cues and a blood meal. (**B**) Percentage of animals fed. Ordinary one-way ANOVA with Tukey’s multiple comparisons test. Columns labeled with different letters are significantly different from each other (P < 0.05). P > 0.05: not significant (ns). Bars and error bars show mean ± SEM. Dots represent individual trials (n = 7–10 trials with 27–31 individuals/trial).

## Genetic disruption of *pdf* alters neuronal morphology

In insects, PDF is normally expressed in ventral clusters of clock neurons that appear during larval development (*49, 50*). Using immunohistochemistry to characterize PDF cell morphology across developmental stages, we find three anatomically and developmentally distinctive groups of PDF+ neurons (**fig. S3A–D**): small ventrolateral neurons (sLNv) appear before the fourth larval (L4) stage, sending projections to the medial brain and the developing optic lobe and later branching to the dorsal brain in late L4 stage. This is distinct from the developmental pattern observed in *Drosophila*, where the small LNv cells and their dorsal projections develop early and persist, largely unchanged, into adulthood (*49*). Around the ventrolateral region, two groups of larger cells (lLNv-d and lLNv-v) begin to appear during late L4 stage that densely arborize the dorsal and ventral halves of the developing medulla (**fig. S3A–E**). The three PDF+ cell groups and their projections persist into adulthood and we observe 8–10 cells in the sLNv group and 4–5 cells in each of the two lLNv groups, consistent with previous studies (*28*). We also noticed soma size differences across all three groups (**Fig. 3A and B**) and differences in PDF staining intensity between small and ventral large cell populations in wildtype animals (**Fig. 3C**). Since homozygous *pdf* mutants completely lose PDF expression (**fig. S1D**), we compared PDF staining between wildtype and heterozygous animals. Although we did not detect differences in overall levels of PDF staining intensity between these genotypes (**Fig. 3C**), we observe morphological differences between wildtype and heterozygous animals. Heterozygous females lose soma size and staining intensity differences between clock cell subgroups and show abnormalities in the dorsal projections from the sLNv cells (**Fig. 3B–F**). These sLNv projections are normally highly stereotyped and form critical connections between the PDF+ cells in the ventral brain and PDF-clock cells in the dorsal brain (*25, 26, 28, 51*–*53*). However, in *pdf* heterozygous animals the projections are defasiculated and fail to reach their normal regions of innervation in the dorsal brain (**Fig. 3D–F**). This phenotype is present during larval development (**fig. S4**) and is similar to deficits observed in *Drosophila pdf* mutants (*54*) and *Drosophila cycle* mutants that also exhibit reduced PDF expression (*55*). Overall, *pdf* heterozygous females show loss of size differences in clock cell bodies, morphological abnormalities, and mistargeting in sLNv projections to the dorsal brain. These data demonstrate that there are anatomically and developmentally distinct groups of PDF-expressing cells in the *Aedes aegypti* brain and that disruption of a single copy of the *pdf* gene leads to specific anatomical deficits in clock cell morphology and projection patterns.

**Figure 3.**
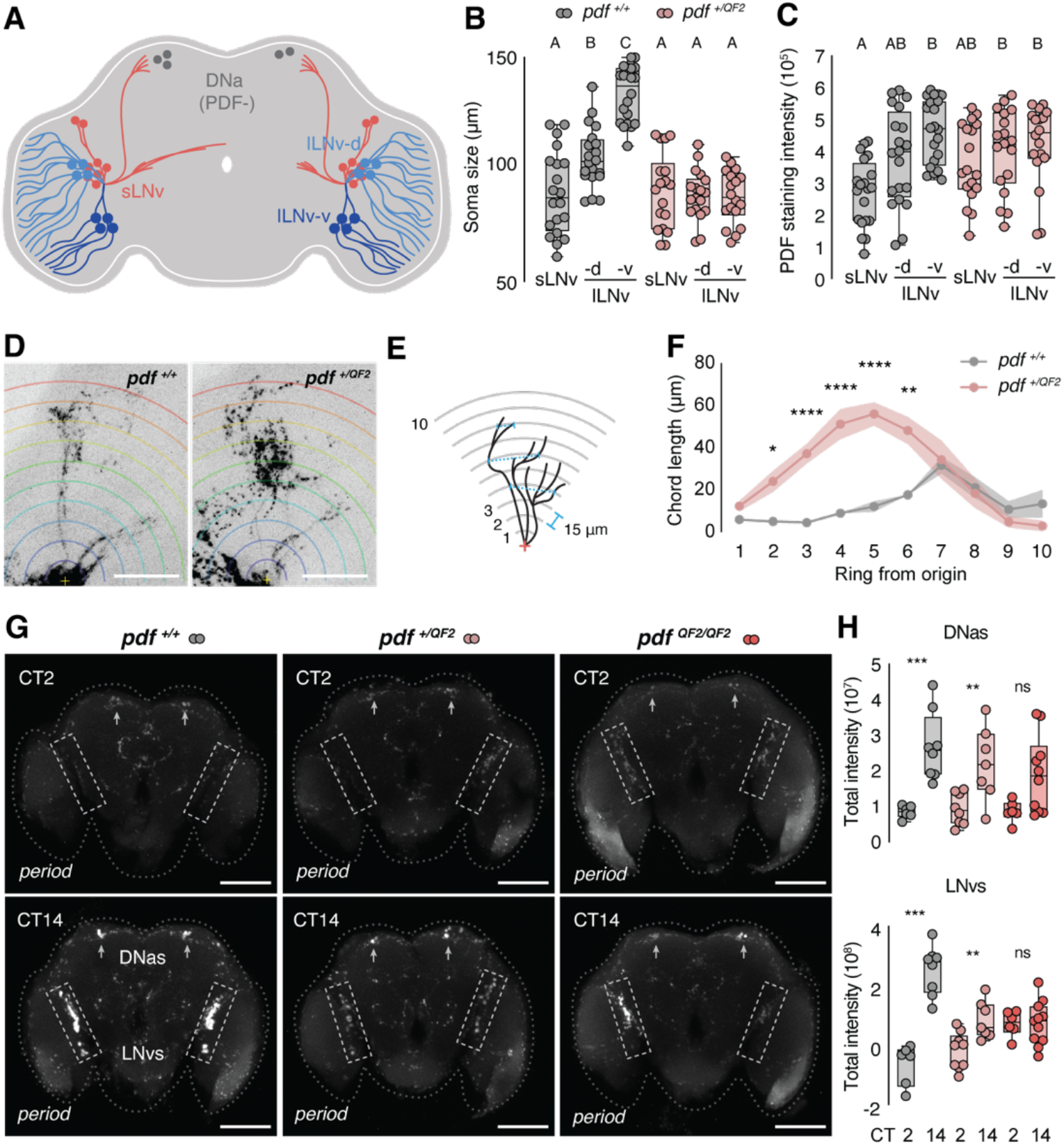
*Pdf* mutation leads to changes in clock cell morphology and disrupted *period* oscillation. (**A**) Schematic of the *Aedes aegypti* circadian neuronal network (sLNv: small ventrolateral neuron, lLNv: large ventrolateral neuron, DNa: anterior dorsal neuron). (**B**) Measurements of PDF neuron soma size (n = 10 for *pdf* ^*+/+*^ and n = 9 for *pdf* ^*+/QF2*^). Each dot represents one soma and two cells per neuron group were measured from each brain. Ordinary one-way ANOVA with Tukey’s multiple comparisons test. Columns labeled with different letters are significantly different from each other (P < 0.05). (**C**) Measurements of average PDF staining intensity (n = 10 for *pdf* ^*+/+*^ and n = 9 for *pdf* ^*+/QF2*^). Each dot represents one soma and two cells per neuron group were measured from each brain. Kruskal-Wallis test with Dunn’s multiple comparisons. Columns labeled with different letters are significantly different from each other (P < 0.05). (**D**) Representative staining of PDF+ neuron dorsal projections. Scale bar: 50 μm. (**E**) Schematic of modified Sholl analysis. Rings were defined from the origin of the dorsal projections in 15 μm increments. (**F**) Chord length measurements from the modified Sholl analysis (n = 8 for *pdf* ^*+/+*^ and n = 10 for *pdf* ^*+/QF2*^). Mann-Whitney test. (**G**) Representative RNA *in situ* hybridization staining of *period* mRNA. Rectangles with dashed line indicate the ventrolateral neurons (LNvs) and arrows indicate the anterior dorsal neurons (DNas). Scale bar: 100 μm. (**H**) RNA *in situ* hybridization staining intensity of *period* mRNA in DNas (top) and LNvs (bottom) (DNas: n = 6–10 for all genotypes per time point; LNvs: n = 5–11 for all genotypes per time point). Mann-Whitney test. Shaded lines show mean ± SEM. Dots represent individual measurements. Boxes extend from the 25th to 75th percentiles. *P < 0.05; **P < 0.01; ***P < 0.001; ****P < 0.0001; P > 0.05: not significant (ns).

## PDF is required for normal *period* transcript oscillation

Previous work in *Drosophila* has demonstrated that PDF synchronizes oscillations in core clock transcripts and proteins in multiple clock cell subgroups (*22, 23*). Apart from the PDF+ ventrolateral neurons, we consistently observed *period* and *cycle* transcript co-localization in an anterior subset of dorsal cells that has previously been reported to show clock protein expression (*28*) (**Fig. 3A**). To ask if loss of PDF changes rhythmic expression of core clock transcript levels, we used *in situ* hybridization to quantify circadian changes in *period* transcript in PDF+ ventrolateral neurons (LNvs) and PDF-anterior dorsal neurons (DNas) (**Fig. 3G**). In wildtype animals, *period* transcript levels are significantly higher at CT 14 compared to CT 2 in both groups of clock cells, similar to previous reports of transcript oscillations (*15, 56*) (**Fig. 3H**). Changes in *period* transcript levels are preserved in heterozygous animals, although amplitude is reduced and these changes are lost in homozygous *pdf* mutants (**Fig. 3H**). Interestingly, we also observed that *period* signal intensity often showed dramatic variation across hemispheres of the same brain in homozygous *pdf* mutants at CT 14 (**Fig. 3G**), suggesting phase disagreement. These data demonstrate that levels of *period* transcript cycle under constant conditions and that cycling is disrupted in multiple clock cell subgroups when *pdf* is disrupted.

## CO_2_-evoked activity and response persistence are modulated by time of day

Our data show that *Aedes aegypti* females are most spontaneously active in the morning and the evening and that blood feeding occurs at both of these timepoints. Although bimodal patterns of morning and evening biting have long been documented in the field, we lack a mechanistic understanding as to how mosquitoes modulate host-cue responsiveness according to the time of day to give rise to observed biting patterns. Previous work has shown that CO_2_ elicits a persistent increase in activity and induces an internal state of predation that allows integration of multi-sensory host cues, which ultimately leads to more effective host-seeking and biting (*34*). However, it is unknown how features of CO_2_ responsiveness may change during the day. To evaluate changes in responses to host-associated cues, we developed the “MozzieDrome” assay, which combines environmental lighting control and videography with automated stimulus delivery (**Fig. 4A**). This allows us to deliver precisely-timed 30 second pulses of 10% CO_2_ throughout the day (**fig. S5A**) and track the movement of 10 individual females over days. Overall response intensity can be represented with velocity changes throughout the recording (**Fig. 4D**), and behavioral responses can be classified in 5-second fragments as “walk” or “flight” using displacement and velocity for each individual (**Fig. 4B and E**). Wildtype *Aedes aegypti* females show robust behavioral responses to CO_2_ during morning (ZT0), midday (ZT6), and evening (ZT12) timepoints but almost no response to a pulse delivered during the night (ZT18) (**Fig. 4C–E**). Since CO_2_ responses are essentially absent during the night (ZT15, 18, and 21) (**fig. S6**), we focused on timepoints during the day to characterize features of these responses.

**Figure 4.**
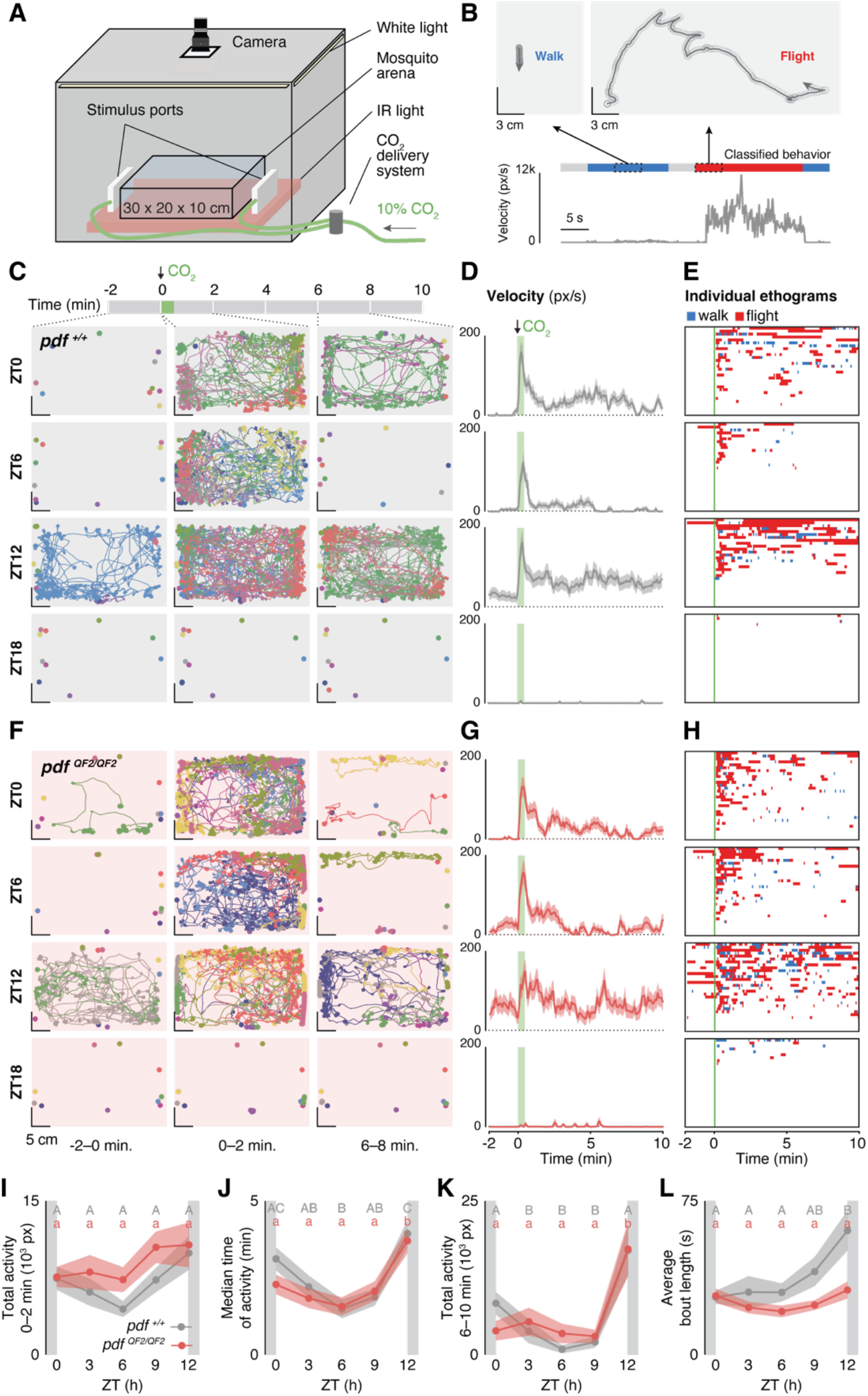
*Pdf* mutants show altered daily modulation in CO2 responsiveness and persistence. (A).Schematic of the MozzieDrome circadian host-seeking assay. (**B**) Example tracking of a mosquito exhibiting walk/flight in two separate 5-second windows (top). Circles denote mosquito positions in every frame. Corresponding velocity plot of a mosquito along with classified behavior in 5-sec windows (bottom). (**C and F**) Representative CO2-evoked activity of groups of 10 *pdf* ^*+/+*^ (C) and *pdf* ^*QF2/QF2*^ (F) mutant females at ZT0, 6, 12, and 18. Activity tracks from selected 2-minute windows in a 12-minute recording from 2 minutes before CO2 stimulus to 10 minutes after the CO2 stimulus. (left: -2–0 min, middle: 0–2 min, right 6–8 min). Track color reflects individual identity and thinner lines reflect higher velocity. (**D and G**) Average velocity of CO2-evoked activity of *pdf* ^*+/+*^ (D) and *pdf* ^*QF2/QF2*^ (G) females at ZT0, 6, 12, and 18 (n = 70–80 for *pdf* ^*+/+*^ and n = 58–59 for *pdf* ^*QF2/QF2*^). (**E and H**) Ethograms of CO2-evoked activities of individual *pdf* ^*+/+*^ (E) and *pdf* ^*QF2/QF2*^ (H) females at ZT0, 6, 12, and 18 (n = 70–80 for *pdf* ^*+/+*^ and n = 58–59 for *pdf* ^*QF2/QF2*^). Each row represents one mosquito in the 12-min recording. For clarity, 30 individuals from each time points were randomly chosen. Behavior classified in 5-sec windows as walk/flight as demonstrated in (B). (**I**) Total activity 0–2 min post-CO2 under LD entrainment in the photophase (n = 47–80 for both genotypes at all time points). Kruskal-Wallis test with Dunn’s multiple comparisons. Columns labeled with different letters are significantly different from each other (P < 0.001). (**J**) Median time of post-CO2 activity under LD entrainment in the photophase (n = 29–46 for all timepoints). Kruskal-Wallis test with Dunn’s multiple comparisons. Columns labeled with different letters are significantly different from each other (P < 0.05). (**K**) Total activity 6–10 min post-CO2 under LD entrainment in the photophase (n = 29–48 for both genotypes at all time points). Kruskal-Wallis test with Dunn’s multiple comparisons. Columns labeled with different letters are significantly different from each other (P < 0.05). (**L**) Average duration of activity bouts under LD entrainment in the photophase (n = 29–46 for both genotypes at all time points). Ordinary one-way ANOVA with Tukey’s multiple comparisons test. Columns labeled with different letters are significantly different from each other (P < 0.05). For panels I–L, uppercase letters show statistical grouping for *pdf* ^*+/+*^, and lowercase letters for *pdf* ^*QF2/QF2*^. Shaded lines show mean ± SEM. White and dark grey shading in **I–J** represent light and dark phases, respectively.

A detailed look at the daytime response profiles of wildtype *Aedes aegypti* females (**Fig. 4D and E)** shows a rapid increase in activity within the first two minutes, and sustained activity after this initial, acute response. Throughout the photophase, acute responses to CO_2_ show no significant differences in response time (**fig. S5B**) or total activity during the first 2 minutes after CO_2_ exposure (**Fig. 4I**). However, we observed that response persistence changes throughout the day: a single pulse of CO_2_ drives >10 minutes of sustained activity in the morning (ZT0) and evening (ZT12) but <6 minutes of activity at midday (ZT6) (**Fig. 4D**). To quantify response duration, we measured the median time of activity during the 10-minute trial for all animals that showed an acute response. We observed that response duration that is highest in the morning (ZT0) and evening (ZT12), with reduced duration at midday (ZT6) (**Fig. 4J**). Consistent with the modulation in median time of activity, sustained activity in the last 4 minutes of the trial (minutes 6–10) is highest at ZT0 and 12 and lower at ZT3–9 (**Fig. 4K**). Additionally, we noted time-of-day variation in individual bout length with longest bouts of activity at ZT12 (**Fig. 4L**), suggesting more persistent behavioral patterns in the evening. These differences are due to changes in flight activity, which is highest at ZT0 and 12, while walking behavior remains relatively constant throughout the day (**fig. S5C**). This is consistent with the hypothesis that CO_2_ is a long-range host cue that induces a “global search” for a host to bite. Taken together, these data show that wildtype *Aedes aegypti* females have unique response profiles to a pulse of CO_2_ depending on the time of day. While showing little to no response to CO_2_ at night, females respond quickly during daytime with the most persistent responses occurring in the morning and evening, coincident with periods of maximal spontaneous locomotor activity, and reported biting.

## *pdf* mutants show altered CO_2_ response persistence profile

Given that CO_2_ responses exhibit nuanced daily profiles in terms of acute reaction and response persistence, we next examined how loss of *pdf* affects features of CO_2_ responsiveness throughout the day (**Fig. 4 F-H**). As observed in wildtype animals, the overall amplitude of acute CO_2_ responses does not significantly differ across the photophase in *pdf* mutants (**Fig. 4I**). To ask if the response persistence profile is disrupted in *pdf* mutants, we analyzed overall activity and participation in the last 4 minutes of the assay (minutes 6–10). This modulation is altered in *pdf* mutants; although sustained responses are high in the evening (ZT12), we observe diminished modulation between morning (ZT0) and midday (ZT6) in median time of activity and sustained activity in minutes 6–10 of the trial (**Fig. 4J and K**). This pattern is similar to the flattened profile of morning locomotor activity observed in **Fig. 1C** and may contribute to reduced blood-feeding efficiency at this time (**Fig. 2B**). Unlike wildtype females, we did not observe changes in activity bout length in *pdf* mutant females during the daytime (**Fig. 4L**), suggesting that although their activity is more persistent in the evening, these bouts are more fragmented compared to wildtype females. Overall, loss of PDF does not alter daily patterns of acute CO_2_ responsiveness, but causes changes the CO_2_ response persistence profile showing reduced persistence in the morning compared to wildtype (**fig. S5D**), flattened morning/midday profile of response persistence, and a loss of bout length modulation in the evening.

## *pdf* mutants show altered circadian persistence profile in DD

Although CO_2_-evoked responses show strong daily oscillation under LD entrainment, we next asked if acute and persistent features of CO_2_ responsiveness are under circadian control and maintained without external zeitgebers. Under constant dark (DD) conditions, wildtype *Aedes aegypti* females show reduced and shortened responses to CO_2_ compared to responses in the photophase (**Fig 5A, see also fig. S7**) and we did not detect significant differences in median time of activity throughout the day (**Fig. 5B**). However, females show variation in total activity in acute (0–2 min) CO_2_ responses (**Fig. 5C**). We note that the peak of acute activity occurs at CT9 by the second day in DD conditions, coincident with the evening locomotor activity peak in DD2, likely due to the relatively short free-running period (see **Fig. 1F and H**). Notably, the acute response profile remains intact in *pdf* mutants, suggesting that acute response to CO_2_ is under endogenous circadian control, but is not dependent on PDF. Since general persistence is reduced under DD in wildtype females, most animals are not active during the last minutes of recording (**fig. S7**). We quantified the total activity in minutes 2–10 after CO_2_ application to characterize sustained responses. Although we did not detect significant circadian variation in sustained responses in either wildtype females or *pdf* mutants (Kruskal-Wallis test, P = 0.13 and P = 0.57, respectively), we observe that in wildtype females sustained activity is highest at CT9, with low sustained activity detected at CT3 and CT12 (**Fig. 5E**), and the peaks and troughs are less pronounced in *pdf* mutants (**Fig. 5F**). We also note changes in bout length; while wildtype females show daily variation in bout length with longest bouts at CT9, this is dampened in *pdf* mutants (**Fig. 5G and H, see also fig. S7**). Heterozygous animals showed an intermediate phenotype in CO_2_-evoked responses under LD conditions but were also impaired under DD conditions, consistent with the phenotypes observed in these animals in spontaneous locomotor activity (**fig. S8**). These data indicate that rhythms in CO_2_ responses are maintained under constant conditions and are likely regulated by the circadian clock. While acute CO_2_ responses are unaffected by loss of *pdf*, these mutants show dampened circadian modulation of bout length. This indicates that acute CO_2_ response and CO_2_ response persistence are controlled by distinct mechanisms and that PDF specifically influences aspects of response persistence.

**Figure 5.**
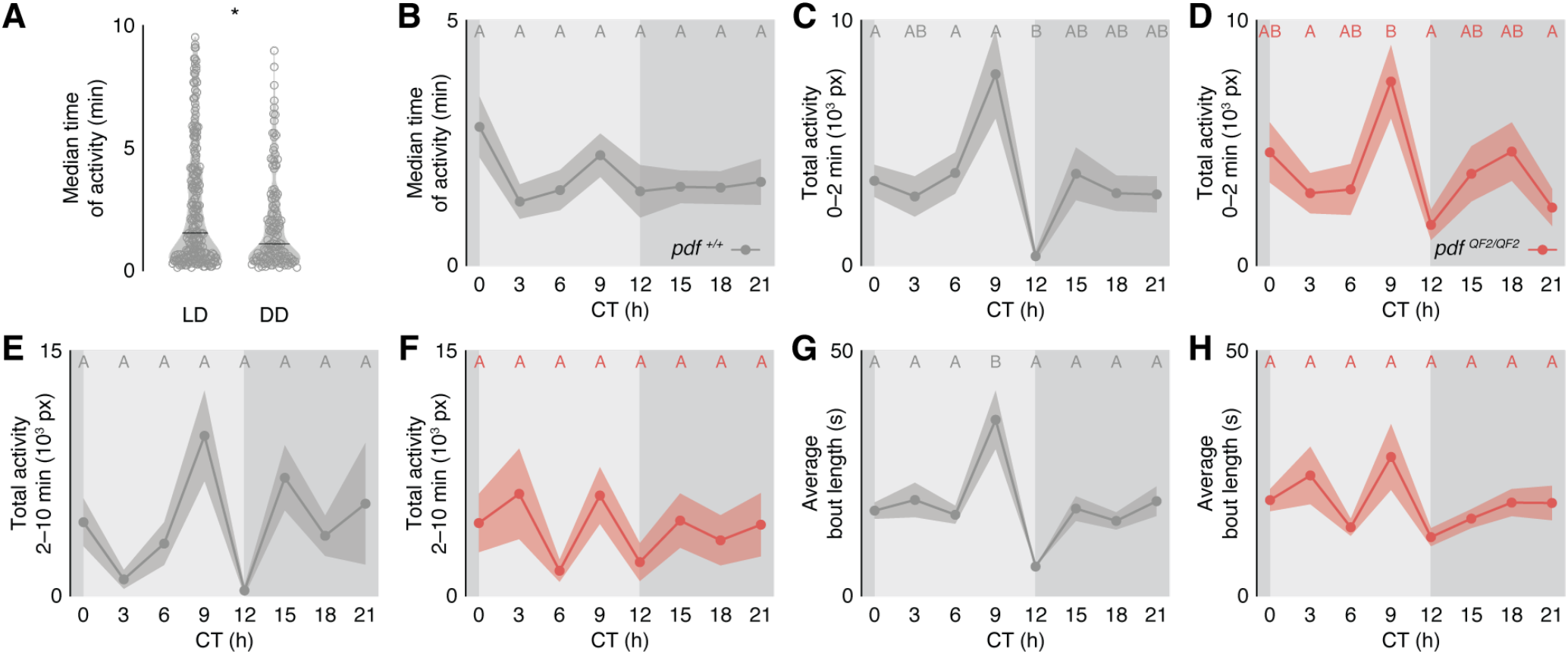
*Pdf* mutants show altered circadian persistence profile. (**A**) Median time of post-CO2 activity of *Aedes aegypti* females under LD and DD entrainments (n = 228 for LD and n = 131 for DD). Mann-Whitney test. *P < 0.05. (**B**) Median time of post-CO2 activity of *Aedes aegypti* females under DD entrainment (n = 7–23 for all timepoints). Kruskal-Wallis test with Dunn’s multiple comparisons. Columns are not significantly different from each other (P > 0.05). (**C and D**) Total initial CO2-evoked activity (0–2 min post-CO2) of *pdf* ^+/+^(C) and *pdf* ^*QF2/QF2*^ (D) females on the second DD cycle (n = 29–30 for both genotypes at all time points). Kruskal-Wallis test with Dunn’s multiple comparisons. Columns labeled with different letters are significantly different from each other (P < 0.05). (**E and F**) Total sustained CO2-evoked activity (2–10 min post-CO2) of *pdf* ^+/+^(E) and *pdf* ^*QF2/QF2*^ (F) females on the second DD cycle (n = 7–25 for both genotypes at all time points). Kruskal-Wallis test with Dunn’s multiple comparisons. Columns are not significantly different from each other (P > 0.05). (**G and H**) Average duration of activity bouts of *pdf* ^+/+^(G) and *pdf* ^*QF2/QF2*^ (H) females on the second DD cycle (n = 7–25 for both genotypes at all time points). Ordinary one-way ANOVA with Tukey’s multiple comparisons test. Columns labeled with different letters are significantly different from each other (P < 0.05). Shaded lines show mean ± SEM. Light and dark grey shading in **B–H** represent respective light and dark phases from prior LD entrainment, respectively.

## Discussion

Although PDF has been implicated in the control of daily locomotor activity in many insects, its behavioral effects vary by species (*24*–*26*). Our data suggest that in *Aedes aegypti*, PDF promotes morning activity, advances morning oscillators, maintains activity after lights-off, and promotes rhythmicity under constant conditions. Our data also suggest that PDF affects molecular oscillations in multiple clock cell subgroups with *pdf* mutants showing altered *period* transcript oscillation in DD. Interestingly, we observe behavioral phenotypes in heterozygous animals, which express one copy of the QF2 transgene and have one intact copy of the *pdf* gene. Although overall levels of PDF are not affected in these animals, we observe anatomical deficits in the dorsal projections of the small LNv cells, which are thought to be critical for communication between ventral populations of PDF+ clock cells and PDF-dorsal clock cells. Our findings suggest that mistargeting of these projections does not result in locomotor activity deficits when environmental timing cues are present, but that deficits are revealed under constant conditions. Although previous studies in *Drosophila* have associated changes in PDF levels with morphological abnormalities and behavioral deficits, the mechanism by which changes in *pdf* dosage contribute to mistargeted projections are of interest for future study (*54*).

*Aedes aegypti* are generally diurnal and we report that females show almost no response to CO_2_ at night, when minimal biting by this species has been reported. Acute responses to CO_2_ are similar across the light phase of the day and this initial response is not affected by the loss of PDF. This is consistent with the hypothesis that the acute CO_2_ response is controlled by peripheral clocks expressed in the maxillary palps that function independently of PDF, similar to the antennal control of olfactory responses in *Drosophila* (*29*). We report for the first time that CO_2_ response persistence is under circadian control and is affected by the loss of PDF. Specifically, PDF reduces morning spontaneous locomotor activity, CO_2_ persistence, and biting efficiency. Overall, we observed unique time-of-day-dependent features of acute and sustained CO_2_ responses that cannot be fully accounted for by the timing of spontaneous activity. Although spontaneous locomotor activity is low at both midday and night timepoints, acute CO_2_ responsiveness remains intact at midday while it is almost completely suppressed at night. We hypothesize that maximal host-seeking efficacy requires the coordination of sustained locomotor activity with chemosensory sensitivity, and blood feeding drive (**Fig. 6**). Daily regulation of sensitivity to internal and external cues may coordinate periods of maximal peripheral sensitivity with internal motivation state to optimize temporal integration of host-associated cues (*57*). Understanding the relationship between sensory coding and locomotor activity to drive appropriate behaviors at specific times of day is a critical area for future research. Our findings suggest that PDF may act on down-stream central brain cells that control overall response persistence (*58*). One area of ongoing work will be to determine whether this behavioral persistence is unique to host-seeking or generalizes to other behaviors.

**Figure 6.**
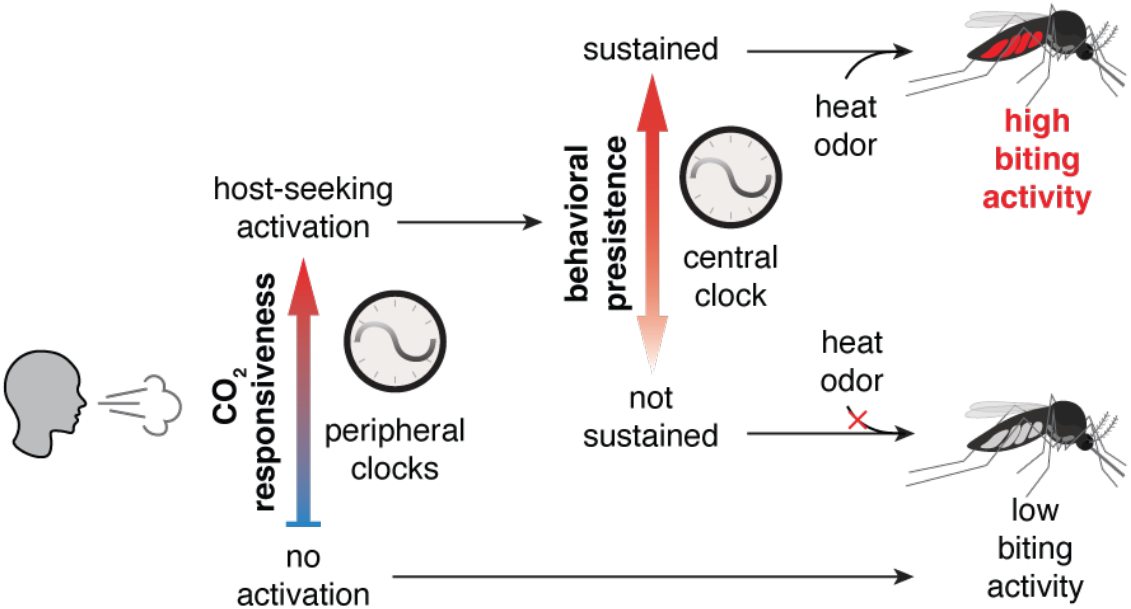
Proposed model of circadian modulation of mosquito biting.

Previous studies have shown that CO_2_ is a potent activator of host-seeking and a single exposure to CO_2_ can drive sustained flight activity and enhance detection and responses to multi-modal host cues (*32, 34, 39, 40, 59*). Our findings indicate that mosquito predation of humans is more effective at specific times of day that are coordinated by central brain clock cells. Understanding the mechanisms of daily rhythms of host-seeking effectiveness can provide new targets to disrupt these and potentially lock females into a “low host-seeking” state. In addition, understanding these changes can identify features of activity that predict biting rhythms and facilitate effective application of current mosquito control technologies by ensuring that they are deployed at the right time of day for maximal efficiency to protect people from disease.

## Acknowledgements

We thank Thomas Gabel for assistance with animal husbandry, Teodora Bratu for assistance in immunohistochemistry staining and image analysis, Nicole Lin for earlier versions of the double-plotted actogram visualization of locomotor activity data, Abhilash Lakshman for developing the phaseR program, and Dan Gross for assistance with Mozziebox construction. We thank Paul Taghert, Darcy Kelley, Trevor Sorrells, Maria de la Paz Fernández, and members of the Duvall lab for comments and useful discussions on the manuscript.

## Author contribution

L.D. carried out or supervised all experiments and data analysis with additional contribution from coauthors. R.H. provided electrical engineering expertise and helped design, construct, and program the MozzieDrome assay. J.M.B. performed image analysis in **Fig. 3** and assisted with the blood-feeding assay in **Fig. 2**. C.G. generated the ILP3 antibody. L.D. and L.B.D. together conceived the study, designed the figures, and wrote the paper with input from all authors.

## MATERIALS AND METHODS

### Mosquito strains

The following strains (*Aedes aegypti* Liverpool background) were used in this study: wildtype *Aedes aegypti* Liverpool, *pdf-QF2* (this study), *QUAS-mCD8-GFP* (*45*).

### Mosquito rearing

*Ae. aegypti* Liverpool mosquitoes and other specified mutant strains were maintained and reared at 28°C, 70–80% relative humidity with a 12 h light: 12 h dark schedule, unless otherwise stated. Eggs were hatched using 1 L of TetraMin fish food suspension in deionized deoxygenated water (one tablet/L). Larvae were fed with TetraMin fish food tablets when needed. Adult female mosquitoes used for experiments were between 3 and 21 days old and co-housed with males, with *ad lib* access to 10% sucrose solution until further experimental procedures. For passaging and mutant generation, adult female mosquitoes were blood-fed on defibrinated sheep blood (Quad5) with 1 mM ATP at 37°C using Hemotek artificial membrane feeders.

### Generation and maintenance of the *pdf-QF2* knock-in knock-out line

Single guide RNAs (sgRNAs) were designed using CRISPR RGEN Tools (http://www.rgenome.net) to target the *Aedes aegypti pigment-dispersing factor* gene (AAEL001754) using the AaegL5 target genome. Efficiency of the sgRNAs were evaluated by injecting sgRNA and Cas9 protein into 200 *Aedes aegypti* Liverpool embryos. The synthesis of the sgRNA and the preparation of an injection mix was described in a previous publication (*44*): DNA oligonucleotide templates were amplified using the KOD polymerase and purified with NucleoSpin columns. T7 Megascript Kit was used for in vitro reverse transcription. RNA was purified using MegaClear column purification kit. Guide RNA (40 ng/μL at final concentration) was purified by ethanol precipitation and mixed with Cas9 protein (300 ng/μL at final concentration). Guide RNA with the most efficient cutting rate (GAAGATGATAGAA-GCACACA**<u>CGG</u>**, PAM sequence underlined) as reflected in frequency of indel mutations was used to design a plasmid for homology-directed repair (HDR). The genetic insert contains the left homology arm (sequence 300 bp before the cut site) and followed by the ribosomal skipping factor T2A, transcriptional activator QF2, eye-specific promotor 3xP3, fluorescent marker dsRed, and the right homology arm (sequence 300 bp directedly following the cut site). This 3169 bp genetic insert was synthesized and cloned into a pUC57 plasmid (GenScript). Plasmid sequence can be found in Data S1. This donor plasmid (700 ng/μL at final concentration) was mixed with guide RNA (40 ng/μL at final concentration) and purified by ethanol precipitation before mixing with Cas9 protein (300 ng/μL at final concentration) and injecting into ∼ 1000 *Aedes aegypti* Liverpool embryos. All embryo injections were performed by the Insect Transformation Facility at the University of Maryland Institute for Biosciences & Biotechnology Research. Injected animals were outcrossed to wildtype virgin male/females in families of < 25 injected animals to screen for germline transmission of the red eye marker under a Leica MZ10 F fluorescence stereomicroscope.

Transgene insertion was characterized by Sanger sequencing which showed complete integration of the genetic insert into the second exon of the *pdf* gene. We detected the presence of a highly repetitive fragment (∼ 2.5 kb) in the intronic region preceding the second exon. However, this did not impact the *pdf*-specific expression of QF2-driven fluorescent marker and the transgene resulted in a null-allele as confirmed by the complete loss of PDF peptide in homozygous *pdf-QF2* mutant (**fig S1**). The *pdf-QF2* mutant line was backcrossed to wildtype Liverpool for five generations before being used for experimentation.

All heterozygous mutants used in this study were generated by continuing backcrossing to wildtype Liverpool animals. Offspring were screened as larvae or pupae by the presence of the red fluorescent eye marker under a Leica MZ10 F fluorescence stereomicroscope. Homozygous animals were established by allowing siblings to interbreed and performing PCR on DNA extracted from single legs of virgin adults using Phire Tissue Direct PCR Master Mix following the manufacturer protocols. A primer pair (F: AGGCACGTCCCAAGTAACAG; R: CGCTGTTGGTGTGGTTAGGA) flanking the insertion were used, and the presence of the long, insertion-containing product and the absence of the short wildtype product was used to identify homozygous individuals (**fig. S1A and C**). Homozygous individuals were pooled to establish lines, which were confirmed by presence of the red eye marker in all offspring when out-crossed to wildtype Liverpool animals. Homozygous lines were maintained for no more than six generations, at which time homozygosing procedures were repeated. We did not observe general mating, blood-feeding, or egg-laying deficits in either the heterozygous or the homozygous *pdf-QF2* mutants.

### MozzieBox locomotor activity assay

The MozzieBox locomotor activity assay was designed after the Flybox (*46*). The overall light-tight enclosure of the MozzieBox is 30 × 30 × 30 cm and is manufactured with expanded PVC sheets. Raspberry Pi computers were used to control light entrainment and image acquisition via a python code. Infrared (IR) LED strips (940 nm) were installed at the bottom of the box, and white LED strips were installed on the top of the box. Two 1/8” translucent acrylic panels were used to provide diffusion of the IR and white light sources. Raspberry Pi 5MP 1080p NoIR cameras were mounted on top of the box and longpass filters (Edmund Optics 43-948) were placed in front of the cameras. Custom printed PCB boards (Digikey) were used to connect IR and white light control to the Raspberry Pi computers. The MozzieBoxes were placed at room temperature, and 27 ± 2°C temperature was maintained throughout.

Mosquitoes were individually loaded into the wells of 12-well tissue culture plates and each well was supplied with 1.5 mL of 10% sucrose in 1% agarose to provide humidity and a food source. Co-housed females were 3–14 days old when loaded into the 12-well plates under cold anesthesia, and loading was performed around ZT10 on the day prior to the first entrainment cycle. Two 12-well plates were placed on the lower diffuser panels to be recorded under programmed light conditions. Three LD cycles, with 30-minute dawn and dusk periods, that were synchronized to the light condition of the main rearing insectary were provided before seven days of DD recordings. Images were taken every minute throughout the recording and later compiled as a 30 frames-per-second video using custom Matlab code (Version 2020b). The videos were analyzed with EthoVision (Version 17) where individual mosquitoes were tracked using gray scaling. Distance traveled between frames were extracted for further analysis. All Mozziebox assays were performed least 3 independent times and genotype, loading position, and the MozzieBox used were randomized between trials. All original codes and additional construction and operational details for the MozzieBox assay are deposited at: https://github.com/DongLinhan/MozzieBox.

### Analysis of locomotor activity

All locomotor activity data were extracted from Ethovision video-tracking outputs using a custom python code. *Pdf*-QF2 animals showed higher levels of lethality at the end of the 10-day trial compared to wildtype and heterozygotes. To ensure that lethality did not influence activity analysis any dead animals at the end of recording were excluded from DD analysis and only animals that survived for more than 72 hours from the end of the last LD cycle were included in LD analysis. Raw distances were transformed into activity scores that represents activity in one-minute bins:

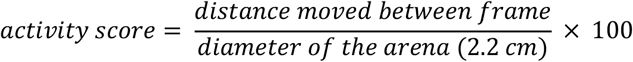

Morning activity phase quantification in **Fig. 1E** was performed with locomotor activity data from ZT22 of the second LD cycle to ZT7 of the third LD cycle. A Savitzky-Golay filter was applied to animal activity scores with a window length of 361 and polynomial order of 3. Peak activity time were extracted from filtered activity score values, and phase was not quantified on animals with low activity (maximum filtered activity score < 5 or total filtered activity score < 100).

Calculation of rhythmic power and period in **Fig. 1G and H** were performed using the phaseR program, developed by Abhilash Lakshman. Chi-square periodogram analysis were applied on the first seven days of DD recordings with the following parameters: alpha for significance tests = 0.01; lowest period = 16; highest period: 32; resolution of period to analyze = 10 mins. Animal is considered rhythmic if rhythmic power > 50. Period is not shown for arrhythmic animals.

Evening activity termination time shown in **fig. S2B** were determined by the first presence of a 5-minute inactivity period from ZT11.5 to ZT13.5. Inactivity is defined as activity score ≤ 2.

Phase quantification shown in **fig. S2C** were performed using custom python code. A Savitzky-Golay filter was applied to animal activity scores in DD1, 2, and 3 with a window length of 361 and polynomial order of 3. Peak activity time were extracted from filtered activity score values within the CT day. Phase was not quantified on animals with low activity (maximum filtered activity score < 5 or total filtered activity score < 100). Phase synchrony was determined using the Rayleigh R test with custom python coding. Raw distances were also binned into 15-minute groups for activity profiles and total activity quantifications shown in **Fig. 1B, C, D and F**. All original code used for locomotor activity data analysis is deposited at https://github.com/DongLinhan/MozzieBox.

### Blood-feeding assay

Co-housed female mosquitoes were entrained for at least 3 days under a 12 h light: 12 h dark schedule and then loaded into a 24.5 × 24.5 × 24.5 cm cage 19–27 hours prior to an experiment in groups of 27–31. The cage was supplied with a cotton wick soaked with DI water to prevent dehydration and kept under the same environmental and light condition until experimentation. Experiments were conducted at ZT0–1 and ZT11–12. Cages were placed inside an environmentally-controlled cabinet (28°C, 75%–85% relative humidity) and acclimated for 10 minutes before the start of an experiment, when a 30-second 25 SCFH pulse of 10% CO_2_ was supplied to the cabinet that elevated CO_2_ level by ∼ 1000 ppm. The cabinet door was opened by 3 cm 2 minutes after CO_2_ application to allow CO_2_ level to return to baseline level. Six minutes after CO_2_ delivery, a Hemotek blood-feeder at 37°C was supplied to the testing cage for 10 minutes, and the cabinet door was closed. The Hemotek blood-feeder was assembled with 3 mL of defibrinated sheep blood and 30 μL of 200 mM ATP. The feeder was covered in a 4 × 4 cm piece of human-scented nylon. Nylon stocking was worn on the forearms of the same experimenter for 6–8 hours then rubbed against the experimenter’s chest and arms prior to collection and storage at -20°C until use. Gloves were used for all mosquito handling in the 24 hours before testing to prevent contamination of host odor from other sources. After each experiment, cages were immediately moved to a cold room to be sorted and crushed on a piece of filter paper to determine blood-feeding status by scoring for the presence of fresh blood.

### Generation of polyclonal guinea pig anti-ILP3 antibody

A custom polyclonal antibody was raised in guinea pig against *Aedes aegypti* ILP3, UniProt: A0EZR6 (ThermoFisher Scientific, Rockford IL). Epitope corresponded to residues 130-148 (RNDLIPPRFRKSPRGIVDE).

### Immunohistochemistry

Unless otherwise specified, all representative immunohistochemistry staining was performed on co-housed female mosquitoes 3–21 days old. The following developmental stages shown in **fig. S3** were determined based on morphology and developmental timeline: L4 early (< 6 hours post-ecdysis, 4 days after hatching); L4 late (> 36 hours post-ecdysis, 5.5–6 days after hatching); pupae (> 16, < 18 hours post-pupation, 7 days after hatching); adult (< 24 hours post-eclosion).

For all adult brain immunohistochemistry, mosquitoes were cold anesthetized on ice and moved to 1 mL of fixative (4% formaldehyde in PBS with 0.25% Triton X-100) and nutated for 3 hours at room temperature. For larval brain dissections, the larval heads were cut off from the thorax and fixed for 1–2 hours at room temperature. For pupal brain dissections, abdomens were removed and the remaining cephalothorax was fixed for 3 hours at room temperature. Tissues were washed five times in wash buffer (PBS with 0.25% Triton X-100) before dissection under a microscope. Lamina were removed to avoid background fluorescence from the eye-specific genetic markers in *pdf-QF2* mutants. Dissected brains were placed in blocking buffer (2% normal goat serum and 4% Triton-X in PBS) at 4°C overnight. Brains were washed three times for 10 minutes each with wash buffer, then placed in primary antibody solution for 1–3 days at 4°C. For brains used for quantitative analysis in **Fig. 3**, the incubation period was 2 days to ensure consistency. Primary antibody solutions were made in antibody buffer (2% normal goat serum and 0.25% Triton-X in PBS). The following antibodies were used: mouse anti-PDF antibody (DSHB) at 1:1000 dilution (1:10,000 dilution in fig. S1E); rat anti-mouse CD8 (Invitrogen) at 1:100 dilution, rabbit anti-GFP (Invitrogen) at 1:1000 dilution; guinea pig anti-ILP3 (this study) at 1:500 dilution. Brains were then washed 5 times for 20 minutes each in wash buffer before overnight incubation at 4°C in secondary antibody solution. Secondary antibody solutions were made in antibody buffer with goat anti-mouse-Alexa647, goat anti-rabbit-Alexa488, goat anti-guinea pig-Alexa488, and goat anti-rat-Alexa488 antibodies (Invitrogen) at 1:500 dilution. Brains were washed for three times for 10 minutes each in wash buffer and then mounted in Vectashield mountant with a 0.2 mm deep spacer. Samples were kept at 4°C before imaging.

### RNA *in situ* hybridization (HCR)

RNA *in situ* hybridization (HCR) were performed using reagents and protocols from Molecular Instruments. Custom probes were designed by Molecular Instruments for the *period* (AAEL008141) and *cycle* (AAEL002049) genes. Animals were loaded into 12-well plates and entrained in the MozzieBox locomotor activity assay for at least four LD cycles (12 hour light: 12 hour dark, 30-minute dawn/dusk transitions) before DD cycles begin. Animals were taken out of the MozzieBox at the corresponding CT time points on the third DD day and quickly cold-anesthetized. Animals were then quickly transferred into fixative solution (4% formaldehyde and 0.25% Triton-X in PBS) under dim red light and nutated at room temperature for 3 hours. Animals were washed five times with nuclease-free wash buffer (NF wash buffer, 0.25% Tween-20 in PBS) before dissection under a microscope. Dissected brains were washed once with NF wash buffer and transferred into 400 μL probe hybridization buffer (Molecular Instruments) and incubated at 37°C for 30 minutes. Brains were then transferred into 250 μL of pre-heated probe hybridization buffer with 4 pmol of custom-made probe and incubated with rotation overnight at 37°C. On the following day, probes were washed 5 times with 400 μL probe wash buffer (Molecular instruments) 10 minutes each at 37°C, and then washed twice, 10 minutes each, with fresh 5x SSCT buffer (5x SSC and 0.1% Tween 20) at room temperature. Brains were then transferred into 400 μL of amplification buffer (Molecular Instruments) and incubated for 10 minutes at room temperature before incubation overnight in 100 μL amplification buffer with 9 pmol snap-cooled hairpins with rotation at room temperature. Brains were washed once for five minutes, twice for 30 minutes, and once at five minutes again before mounting in SlowFade Diamond mountant with a 0.2 mm spacer. Samples were kept at 4°C before imaging.

### Brain imaging

All images shown and analyzed were obtained with a Zeiss LSM880 laser scanning confocal microscope unless otherwise noted. All images used for quantitative analysis and representative images in **Fig. 3 B–G, Fig. 3H**, and **fig. S1D** were obtained with a 20x objective. Images used for same cohort of quantitative analysis (PDF intensity quantification in **Fig. 3C** and *period* intensity quantification in **Fig. 3H**) were acquired at 1024 × 1024 pixels and stored as 16-bit images under same imaging parameters and laser setups. Entire brains were captured with 2.3 μm z-step. Epifluorescent imaging was used for representative images shown in **fig. S1E** with a Nikon Ti2E microscope with a 20x objective. Images were acquired with 0.9 μm z-step, at 3892 × 3884 pixels, and stored as 16-bit images.

### Image processing and analysis

All image processing and quantitative analysis were performed with Fiji (version 2.14.0). All representative images of the same cohort were displayed without varying brightness/contrast within the same channel. Composite images shown in **fig. S1D** were maximum projections of slices showing relative PDF/ILP3 signals to minimize background fluorescence from the eye-specific genetic markers in *pdf-QF2* mutants. All other representative images were maximum projections of relevant slices to avoid non-specific fluorescence background. Genotypes were not blinded to the experimenter. All image analysis were performed on the right hemisphere unless brain tissue appeared to be damaged, in which case we use the left hemisphere.

For quantification of PDF staining intensity and soma size, two somas of each PDF+ neuronal group were selected as ROIs and reported in **Fig. 3B and C**. Two somas were sampled from each PDF+ neuron group. For each soma, a single Z-slice with highest fluorescent signal were chosen for defining ROI and measuring average pixel intensity. Background pixel intensity was measured on the same Z-slice. PDF staining intensity was calculated as:

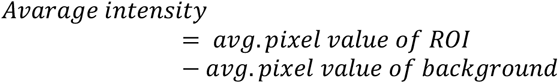

Modified Sholl analysis shown in **Fig. 3E and F** were performed using the SNT toolkit on Fiji. PDF immunostaining images were transformed into a maximum intensity projection of all z-steps and thresholding was performed to highlight the morphology of the dorsal projections. Using the Sholl analysis function on the SNT toolkit, origin was defined as the branching point between the dorsal projection and the medial projection from the small ventrolateral neurons. Sets of 10 rings were drawn surrounding the origin by giving the following inputs: starting radius = 0 μm, increases = 15 μm, final radius = 150 μm, samples per radius = 3. Distance measurements were performed to record the largest chord length defined by the interceptions between staining signal and the rings. Chord length was recorded as zero if no interception was found.

For quantification of RNA *in situ* hybridization staining intensity of *period* mRNA in the ventral lateral neurons, image slices were transformed into 32-bit sum intensity images. To capture all staining signal from the PDF+ ventral lateral neurons, we defined a rotated rectangle at a fixed width of 50 μm that spanned the separation between the central brain and the optic lobe. Four additional ROIs were chosen surrounding the ventral lateral neurons to derive average background intensity. For the anterior dorsal neurons, *period* staining was visible for brains across genotype and circadian timing. Therefore, 10 Z-stacks containing the anterior dorsal neurons were combined to a 32-bit images by applying the sum intensity function on Fiji. An ROI was drawn to include all anterior dorsal neurons in the sum image while minimizing space for background. Four ROIs were chosen surrounding the anterior dorsal neurons to derive average background intensity. Total pixel intensity was calculated as:

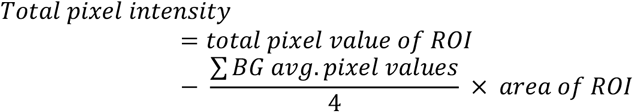

For **fig. S3D**, PDF+ cells were manually counted using confocal images. To ensure overlapping soma signals does not influence cell count accuracy, the hemisphere with less overlapping staining signal and therefore more anatomically defined somas was selected for counting. Cell sizes measurements shown in **fig. S3E** includes all detectable soma with PDF immunoreactivity from both hemispheres, unless presence of dissection/imaging errors was identified. Soma size was measured with FIJI by manually drawing a polygon ROI selection around PDF+ cell body.

### MozzieDrome circadian host-seeking assay

The MozzieDrome host-seeking assay features a circadian entrainment enclosure, a video-recording platform, a mosquito arena with CO_2_ delivery components. Acrylic sheets for all custom-built components were processed with a laser cutter and/or a hand drill. Construction notes, original codes, and vector files used for laser-cutting can be found at: https://github.com/DongLinhan/MozzieDrome. The enclosure of the MozzieDrome was constructed with 1/4’’ black acrylic sheets. A 60 × 45 × 45 cm box with ventilation holes and a 60 × 45 cm hinged door was assembled with acrylic cement. Black foam tape was applied to the rims and secured with duct tape and black-out curtains were installed to ensure a light-proof seal. Four rubber feet were attached to the bottom of the box to provide elevation and soften vibration. Ventilation holes at the top were equipped with two USB cooling fans covered with a light-proof screen and ventilation holes at the bottom were covered with an elevated black acrylic platform to ensure darkness. The lighting system uses white LED strips covered with a 1/8’’ white translucent acrylic sheet to provide diffusion. The white LED light and the CO_2_ delivery system were controlled with Matlab code to provide animals with LD (12 h light: 12 h dark, with 30 min transitions) and DD conditions.

A camera mount was installed on a pre-cut camera hole on top of the enclosure and an IR light platform was placed inside of the enclosure to provide illumination for videography. An IR longpass filter was secured inside the enclosure to block visible light and prevent visible light interfering with video acquisition. Back and side IR illumination was provided with a custom-built light box constructed with 1/4’’ clear acrylic sheets covered with two sheets of Kimwipe to provide diffusion.

The 10 × 20 × 30 cm mosquito arena was constructed with 1/8’’ clear acrylic sheets and secured with acrylic cement. Two pre-cut holes 13 mm in diameter were designed to accommodate sugar feeders. The open faces were covered with mesh sheets secured with hot glue.

CO_2_ delivery was controlled by solenoid valves used to control the flow speed of CO_2_ through 1/4’’ OD tubing and 1/8’’ ID tubing by adjusting the duration of an on/off cycle according to readings from a flow meter. Custom mixed 10% CO_2_ was supplied to the solenoid valve and a T-splitter was used to divert the CO_2_ stream into two diffuser pads (Flystuff).

To avoid contamination with human odor, all elements inside of the enclosure were wiped with isopropanol and handling was done with gloves. The arena was cleaned with deionized water and 70% ethanol between trials. Co-housed female mosquitoes (n = 9–10) reared under the same LD entrainment schedule as in the MozzieDrome were aspirated into the mosquito arena via the sugar feeder hole. Two sugar feeders with 5% sucrose solution were mounted on sugar feeder holders. IR light sources and ventilation fans were turned on at this point and stayed on throughout each experiment. All preparation for each experiment was done before ZT12 of the day prior to the testing period to allow animals to acclimate. For each experiment, testing consisted of 4 days of trials that are 3 hours apart. Days 1 and 2 were under LD entrainment (12 hour light: 12 hour dark, 30 minute transition) and days 3 and 4 were under DD conditions. For each trial, the camera recorded a 12 minute 60 frames-per-second video at the beginning of each testing ZT/CT hour. A 30-second supply of CO_2_ at a flow rate of 7 SCFH was programmed to start at minute two of the corresponding ZT/CT testing hour. Scheduling of video-recording was done with the Window Task Scheduler to call a start/end recording function from the StreamPix software. The StreamPix software was pre-configured with exposure time, gain, acquisition speed, and ROI. Light and CO_2_ delivery were programmed via custom Matlab program. Animals were cold-anesthetized and removed from the mosquito arena at the end of each experiment. No more than one death was recorded at the end of the recording.

### SLEAP animal behavior tracking

Raw videos acquired with StreamPix were batch-processed with the Batch Processing Module to be converted into AVI format. AVI files were then compressed with the command-line software ffmpeg to MP4 format to reduce file size. SLEAP was used to track animal activity and identity. Videos recorded at ZT0 and/or 12 were chosen to train models on SLEAP and multiple trainings were run until all videos were tracked with > 99% of the frames showing accurate tracking. Trained models were applied on datasets of 4–30 videos. Videos of low tracking quality were pooled again for the training of a new model, followed by visual inspections. After all videos were confirmed to be tracked with >99% accurate tracking, all were visually inspected and manually corrected to ensure correct identity assignment in cases of collisions and rare occurrences of other errors (track point on dark spots of the arena, missing track point from blurry shadows of fast flight activity, etc.). Output H5 files were used for further analysis with custom python codes.

### Analysis of CO_2_-evoked response

All data handling for the MozzieDrome assay and all analysis performed for CO_2_-evoked response, unless otherwise noted, were conducted with custom python scripts deposited at https://github.com/DongLinhan/MozzieDrome. Processing of raw H5 files follows analysis guidelines outlined by authors of SLEAP (https://sleap.ai/notebooks/Analysis_examples.html) to fill in missing values, without further processing. For track visualization purposes (**Fig. 4C and F**), animal tracking coordinates were processed with a Savitzky-Golay filter with a window size of 25 frames and a polynomial order of 3. For heatmaps shown in **Fig. 4E and H**, 30 individuals were chosen randomly: pandas.DataFrame.sample (n=30, axis=1, random_state=42). For response time calculation, moving distances between frames were processed with a Savitzky-Golay filter with a window size of 30 frames and a polynomial order of 3. Since the analysis reports behavioral response to CO_2_, inactive animals (total activity in 12 minutes of recording < 500 pixels) were excluded from this analysis. Rolling windows were used to compare total activity in 15 frames before a time point (pre_movement) with the total activity in the 15 frames following the same time point (post_movement). Response incidence was recorded if post_movement > 3 x pre_movement and post_movement > 100 pixels. Response time is defined as the earliest response incidence within 60 seconds of the start of CO_2_ delivery, on animals not showing response incidences 30 seconds before CO_2_ delivery. For behavior classification, tracking results were segmented into 5-second intervals. For each interval, three variables were extracted: vector length (displacement) computed from the starting and ending coordinates, total distance moved, and maximum velocity. Flight was assigned to the interval if any of the follow criteria were met: 1) total distance moved ≥ 175 pixels, or 2) total distance moved < 175 pixels and maximum velocity > 4 pixels/frame. Walk was assigned to the interval if vector length > 5 pixels, and total distance moved < 175 pixels, and maximum velocity < 4 pixels/frame. Animals were considered active if any walk/flight behavior was identified in the given time frame. Intervals were labeled inactive if it did not meet the criteria for walk or flight. For quantification of median time of activity, timestamps post-CO_2_ with flight or walk activity are grouped and median is calculated from all timestamps showing activity. All analysis to quantity post-CO_2_ responses, including quantification of total activity, active time, or bout length excludes individuals that showed walk or flight activity in the in the (−2–0 min) window before CO_2_ application to prevent inclusion of CO_2_-independent spontaneous activity. All analysis to specifically characterize sustained aspects of CO_2_ response further excludes individuals that did not show activity in the first (0–2 min) window post-CO_2_ application. Since CO_2_ responses are generally less persistent under DD conditions, summation of sustained activity (2–10 min) excludes any activity observed after a period of inactivity > 4 minutes to prevent the inclusion of spontaneous non-CO_2_-induced activity.

### Statistical analysis

All statistical analyses were performed with GraphPad Prism or custom python code. Data are shown as mean ± SEM unless noted otherwise. Statistical methods and sample sizes are included in figure captions. Nonparametric tests were used for data that does not follow normal distribution. Data used for all figures, except **fig. S6** and **fig. S7**, along with statistical test details, including exact p-values, can be found in **Data S1**. Data for behavioral classification shown in fig. S6 and fig. S7 can be found in **Data S2**.

**Figure S1.**
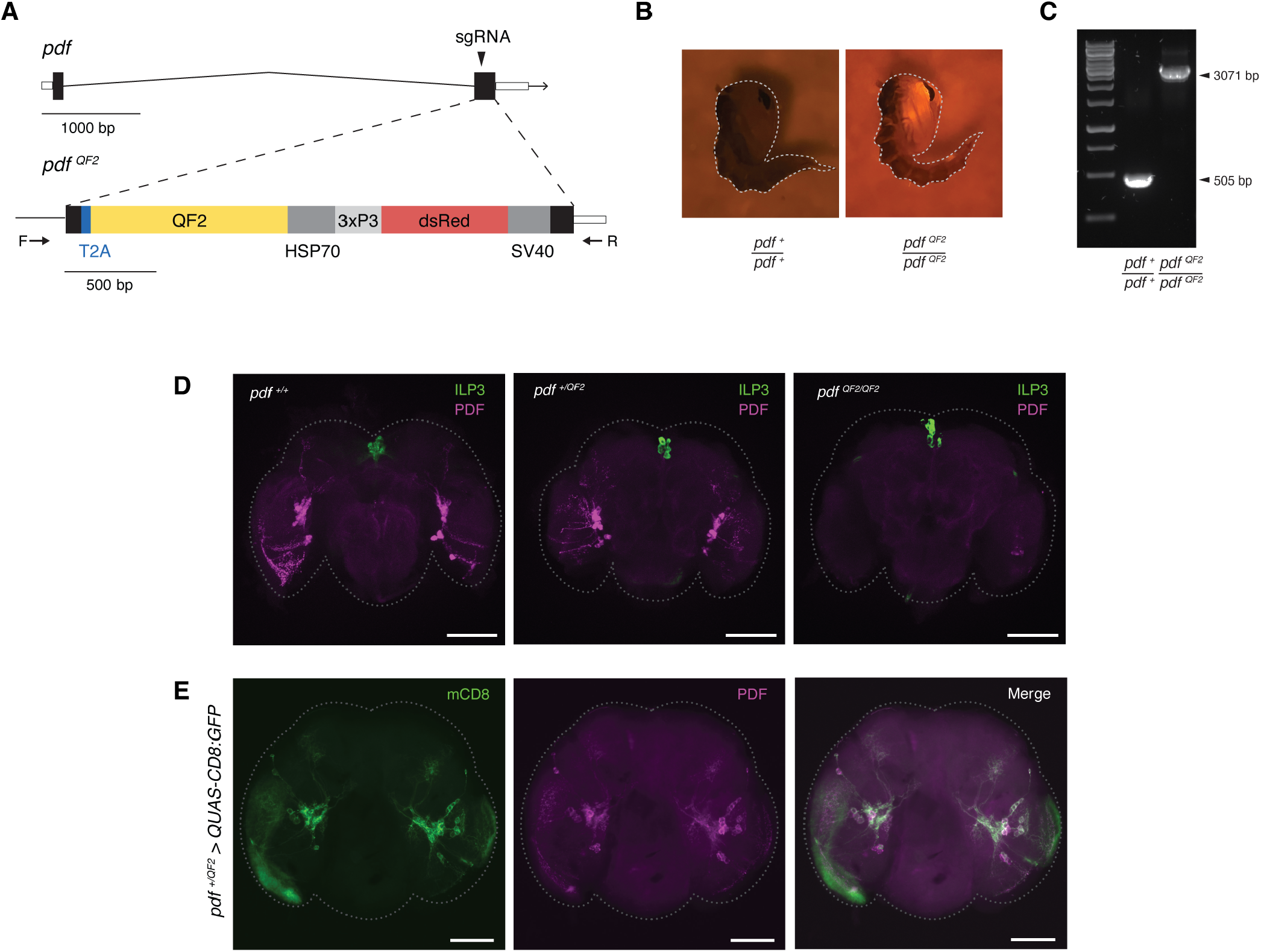
Generation and characterization of the *pdf-QF2* knock-in knock-out mutants. (**A**) Schematic of the genetic insertion using the CRISPR/Cas9 system and homology directed repair. (**B**) Expression of the eye-specific fluorescent marker in female pupae. (**C**) Agarose gel electrophoresis of genotyping product using primers (F and R) labeled in (A). (**D**) Representative images of PDF and ILP3 immunohistochemistry staining on adult female mosquitoes. Scale bar: 100 μm. (**E**) Representative images of mCD8 and PDF staining on *pdf-QF2 > QUAS-mCD8-GFP* adult female mosquitoes. Scale bar: 100 μm.

**Figure S2.**
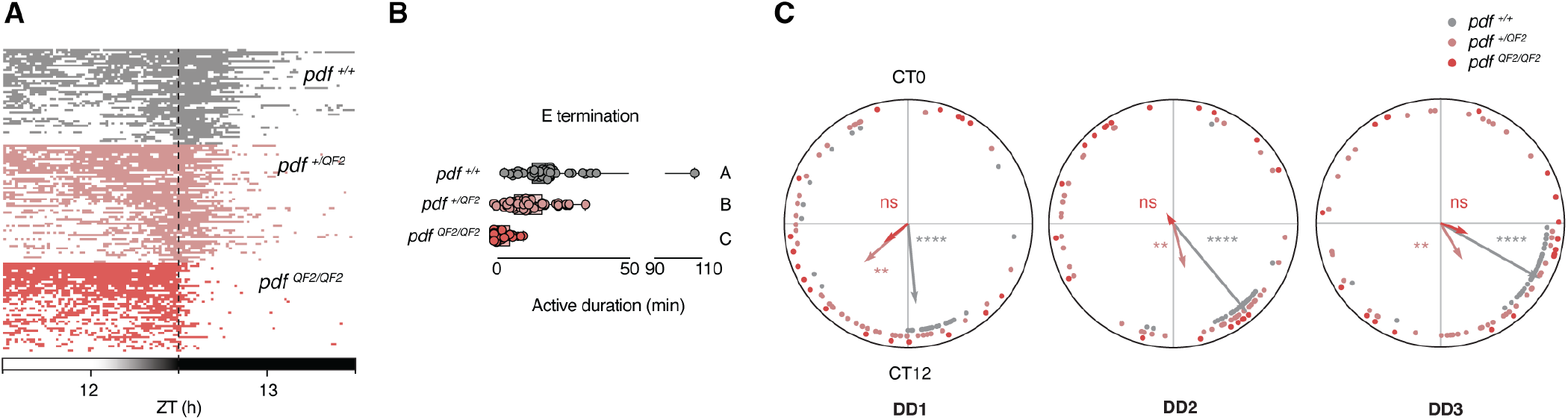
Additional analysis of locomotor activity under LD and DD conditions, related to Fig. 1. (**A**) Evening activity of adult female mosquitoes in Fig. 1C. Activity is shown as colored block if activity score > 2 (see methods). For A and B, n = 43 for *pdf* ^*+/+*^, n = 53 for *pdf* ^*+/QF2*^, and n = 40 for *pdf* ^*QF2/QF2*^. (**B**) Time of evening activity termination (> 5 minutes of inactivity). Kruskal-Wallis test with Dunn’s multiple comparisons. Columns labeled with different letters are significantly different from each other (P < 0.05). (**C**) Circular plots of the phase of activities in the first three DD days (n = 38–40 for *pdf* ^*+/+*^, n = 41–46 for *pdf* ^*+/QF2*^, and n = 18–20 for *pdf* ^*QF2/QF2*^). The vectors (arrows) pointing from the center towards the perimeter show the average peak activity time and vector length corresponds to the extent of phase coherence. Asterisks correspond to the P-value of Rayleigh R test for phase coherence. **P < 0.01; ****P < 0.0001; P > 0.05: not significant (ns). Rows in (A) and dots in (B) and (C) represent individual animals. Boxes extend from the 25th to 75th percentiles.

**Figure S3.**
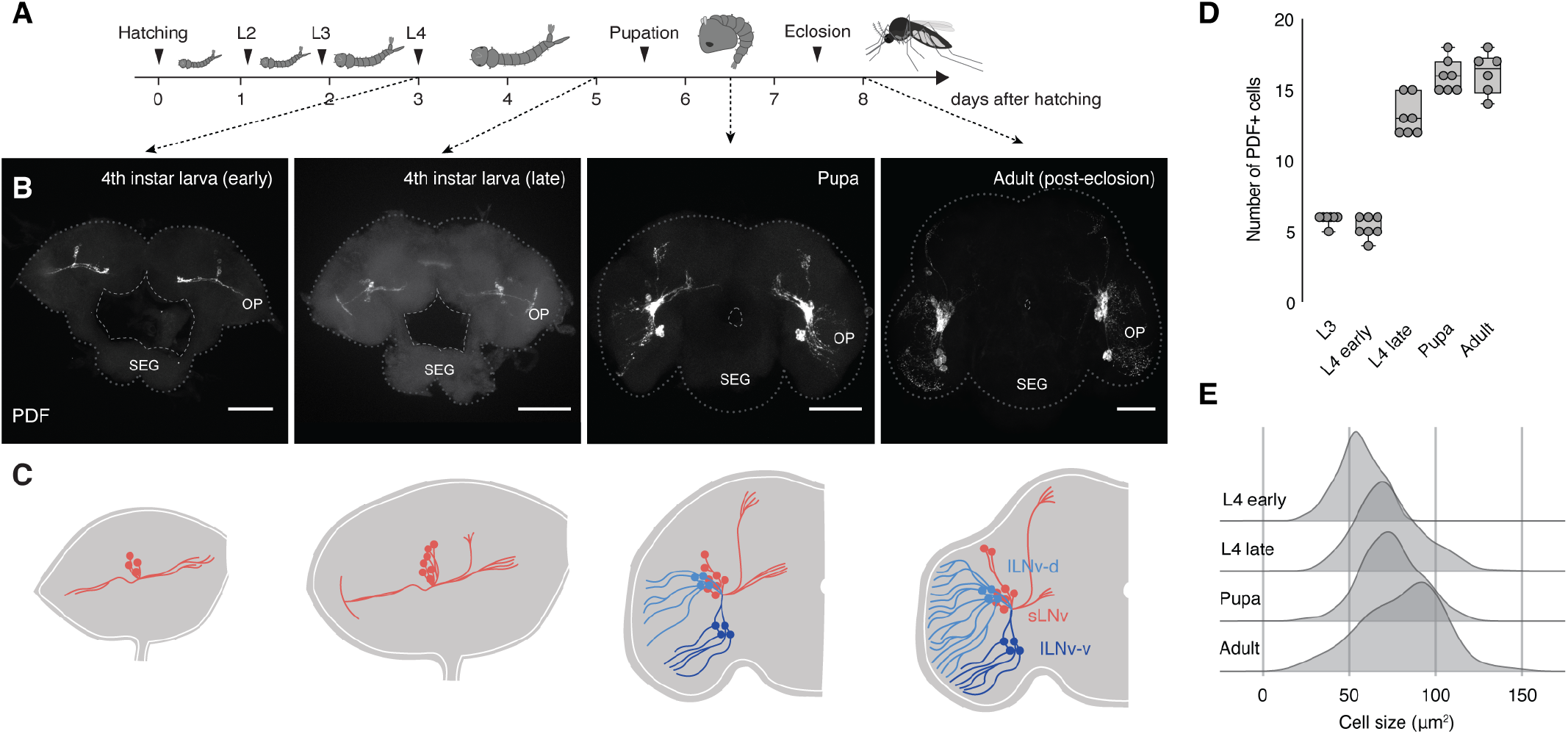
PDF+ neuron development in *Aedes aegypti* mosquitoes, related to Fig. 3A. (**A**) *Aedes aegypti* mosquito developmental timeline, adapted from (*60*). (**B**) Representative PDF staining of the developing brain. OP: (developing) optic lobe; SEG: suboesophageal ganglion. Scale bar: 50 μm. (**C**) Cartoon representation of PDF+ neuronal development. (**D**) Number of PDF-immunoreactive cells across developmental stages (n = 6–7 for each developmental stage). (**E**) Cell size measurements of PDF+ neurons across developmental stages (n = 9, 13, 9, and 5 for L4 early, L4 late, pupa, and adult, respectively).

**Figure S4.**
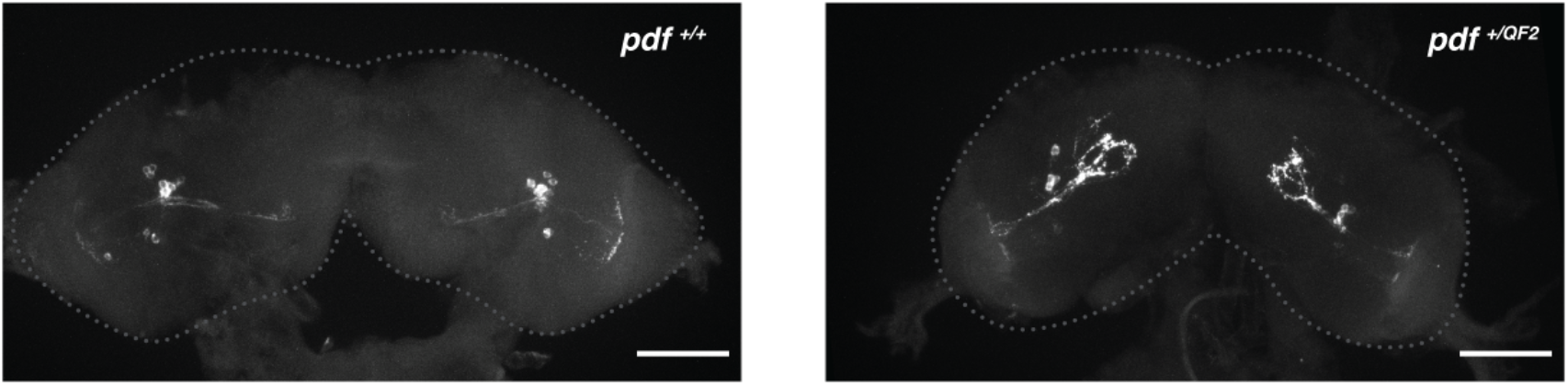
Heterozygous *pdf* mutants show defects in PDF+ neuronal development in *Aedes aegypti* mosquitoes, related to Fig. 3. PDF staining on brains (cerebral lobes) of late stage fourth instar larvae, Scale bar: 50 μm.

**Figure S5.**
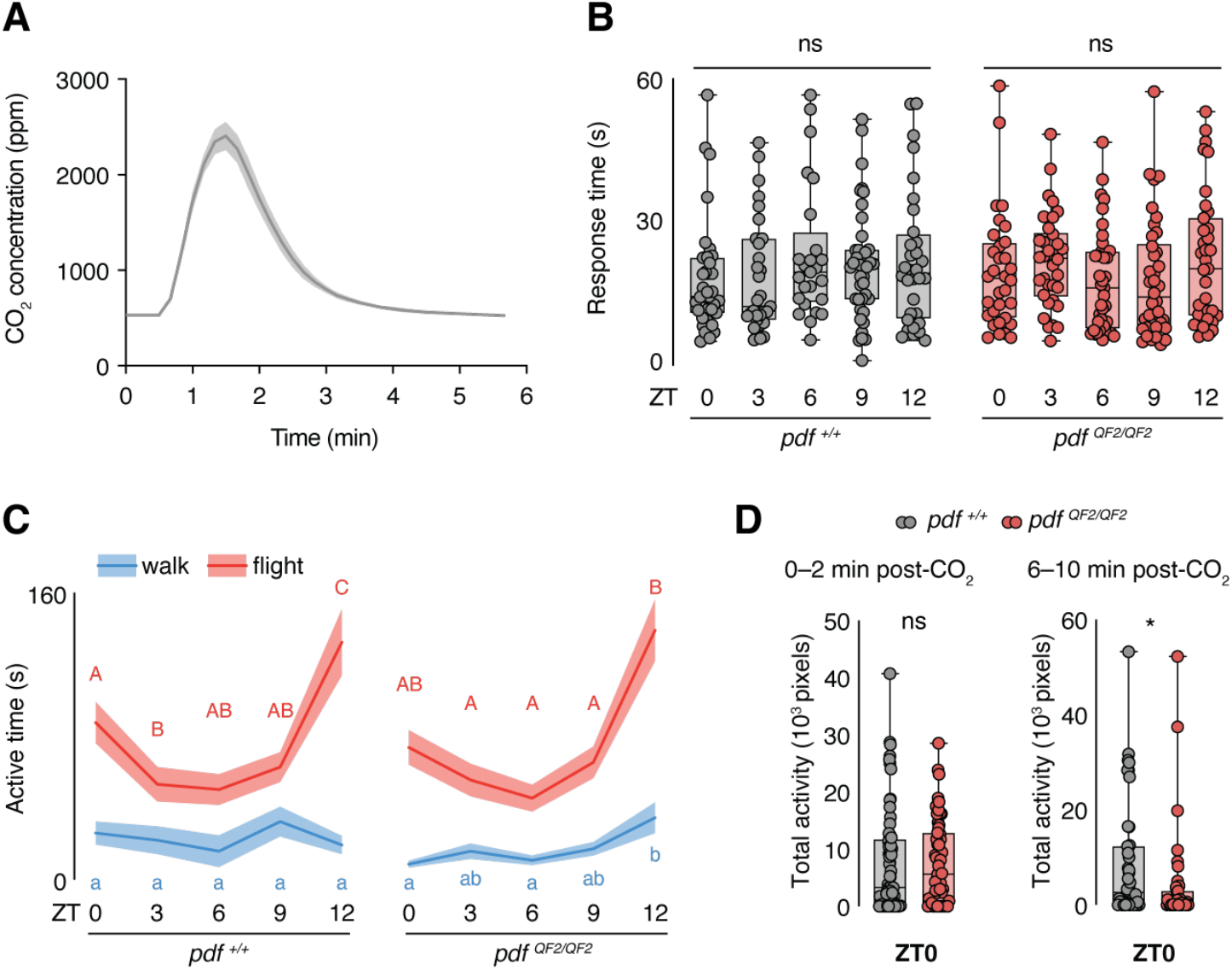
Additional analysis of CO2-evoked responses, related to Fig. 4. (**A**) CO2 concentration measurements in the MozzieDrome assay. Three independent measurements were taken starting from the 30-second 10% CO2 pulse as shown in **Fig. 4A**. (**B**) Activity onset time of *pdf* ^*+/+*^ and *pdf* ^*QF2/QF2*^ females in the photophase. Individuals that were inactive in the 30 seconds pre-CO2 but displayed activity in the 1-minute post-CO2 stimuli were included (n = 25–40 for *pdf* ^*+/+*^ and n = 32–38 for *pdf* ^*QF2/QF2*^). Ordinary one-way ANOVA with Tukey’s multiple comparisons test. (**C**) Total duration of walk/flight behavior post-CO2 in the photophase. Individuals that displayed activity post-CO2 stimuli were included (n = 34–46 for *pdf* ^*+/+*^ and n = 29–42 for *pdf* ^*QF2/QF2*^). Kruskal-Wallis test with Dunn’s multiple comparisons. Data labeled with different letters are significantly different from each other (P < 0.05). Uppercase letters show statistical grouping for flight activity, and lowercase letters for walk activity. (**D**) Total activity 0–2 min (left) and 6–10 min (right) post-CO2 under LD entrainment in the morning (ZT0) (0–2: n = 65 for *pdf* ^*+/+*^ and n = 55 for *pdf* ^*QF2/QF2*^; 6–10: n = 44 for *pdf* ^*+/+*^ and n = 42 for *pdf* ^*QF2/QF2*^). Mann-Whitney test. Shaded lines show mean ± SEM. Dots represent individual animals. Boxes extend from the 25th to 75th percentiles. *P < 0.05; P > 0.05: not significant (ns).

**Figure S6.**
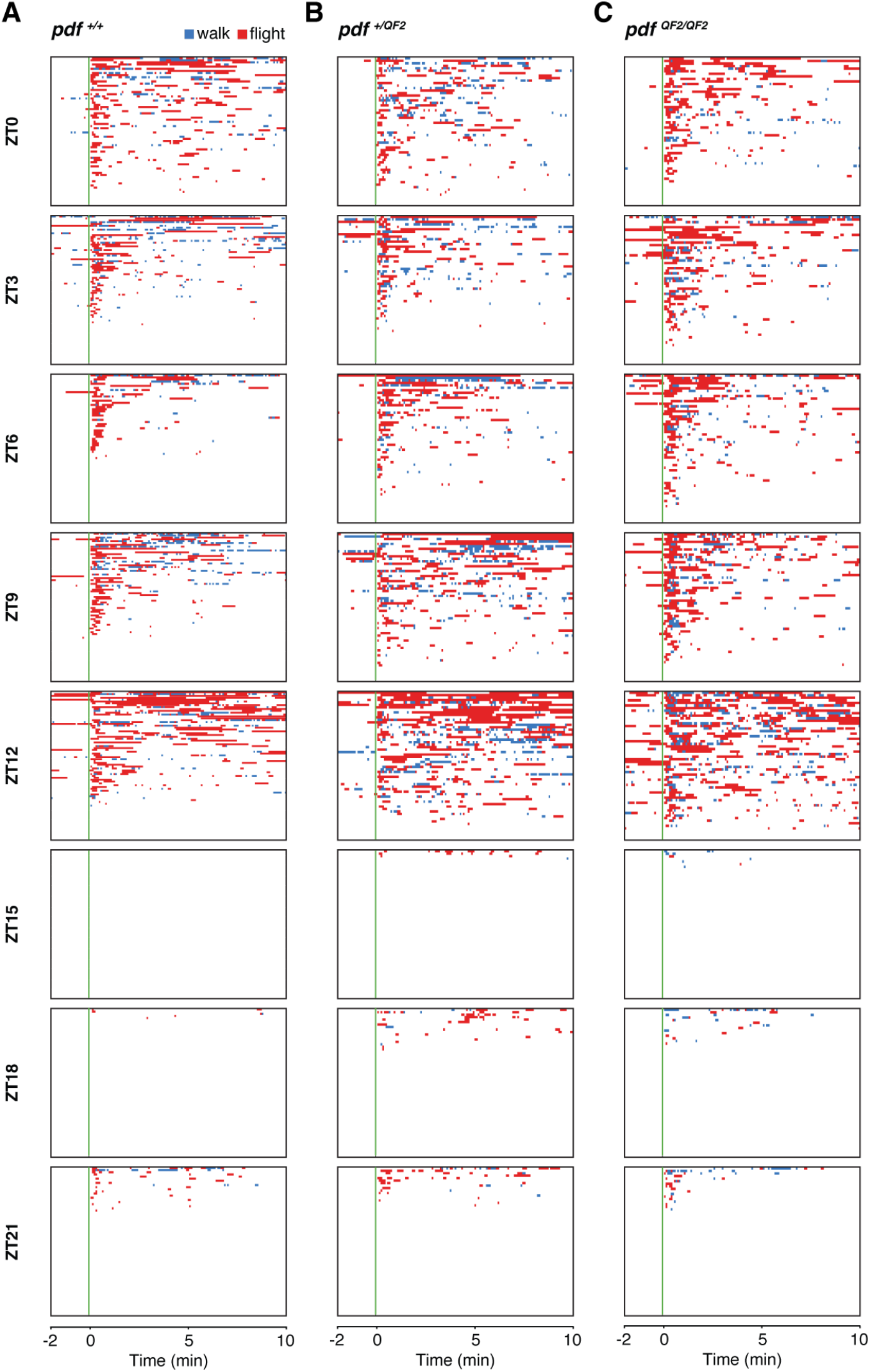
Ethograms of CO2-evoked activities of *pdf* ^*+/+*^ (A), *pdf* ^*+/QF2*^ (B), and *pdf* ^*QF2/QF2*^ (C) mosquitoes under LD entrainment, related to Fig. 4. Rows represent individual animals; n = 59–80 individual mosquitoes for all genotypes at all time points.

**Figure S7.**
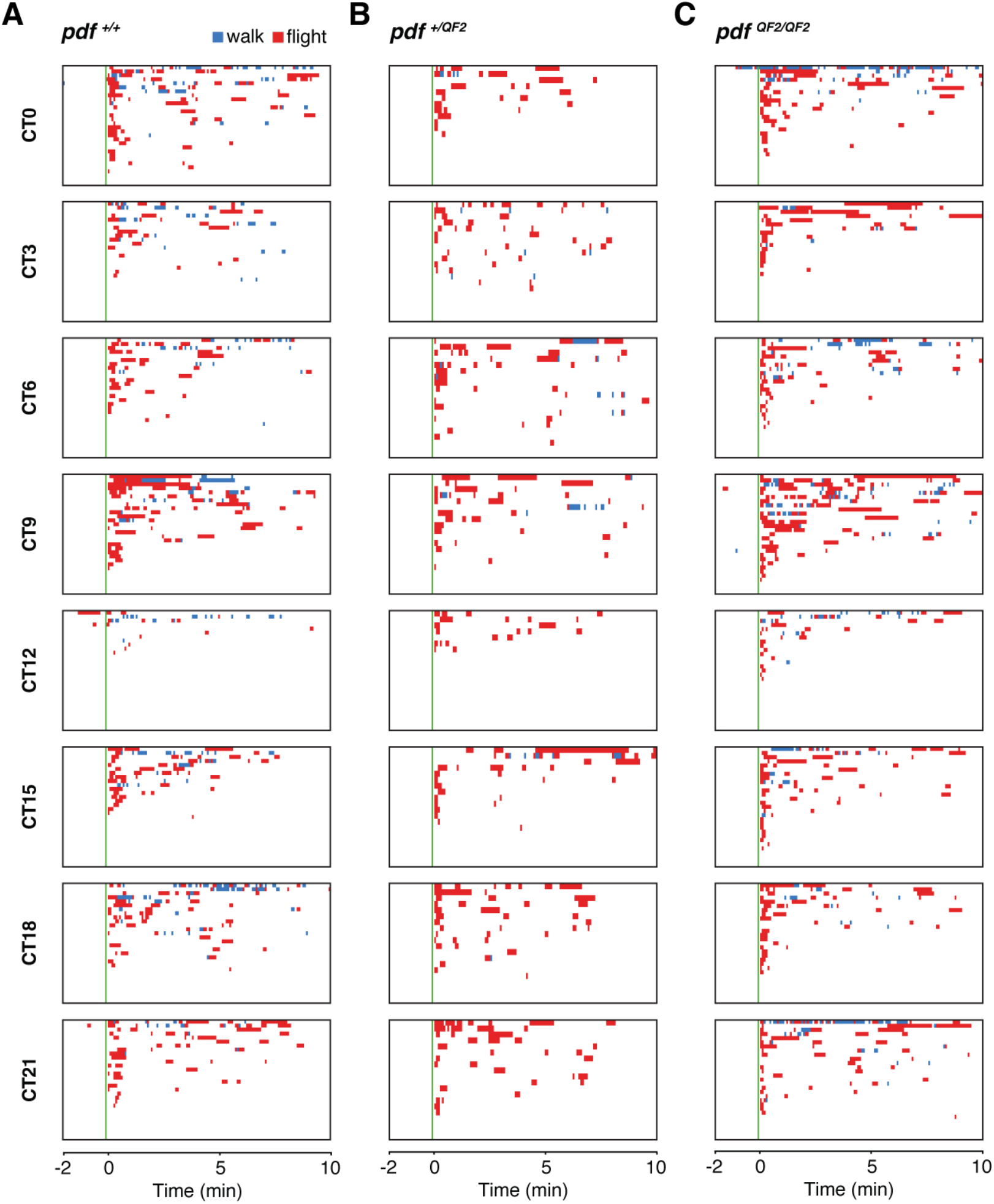
Ethograms of CO2-evoked activities of *pdf* ^*+/+*^ (A), *pdf* ^*+/QF2*^ (B), and *pdf* ^*QF2/QF2*^ (C) mosquitoes under DD entrainment, related to Fig. 5. Rows represent individual animals; n = 20–30 individual mosquitoes for all genotypes at all time points.

**Figure S8.**
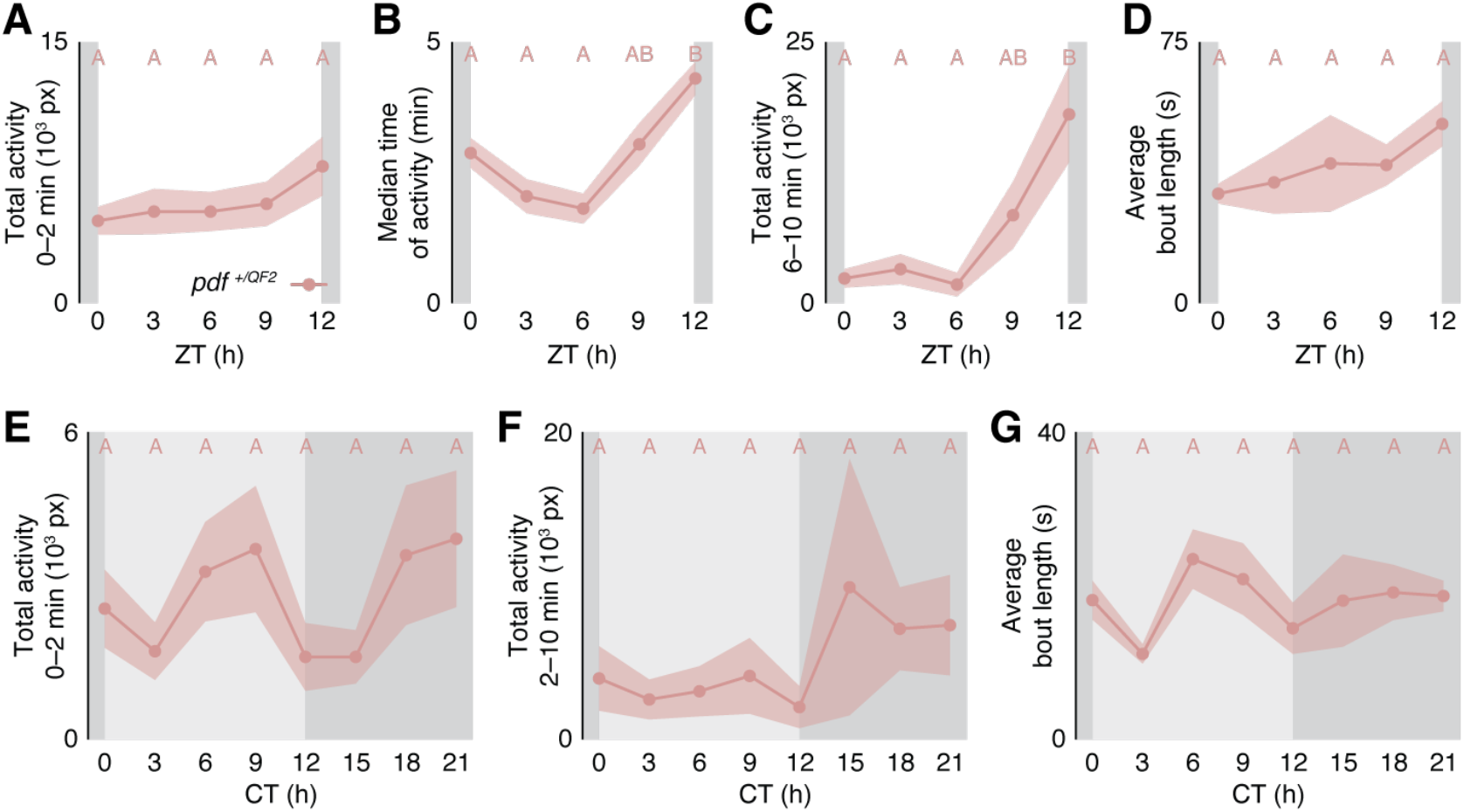
Heterozygous *Pdf* mutants show defects in CO2-evoked response under LD and DD conditions, related to Fig. 4 and 5. (**A**) Total activity 0–2 min post-CO2 under LD entrainment in the photophase (n = 48–60 for all time points). Kruskal-Wallis test with Dunn’s multiple comparisons. Columns are not significantly different from each other (P > 0.05). (**B**) Median time of post-CO2 activity under LD entrainment (n = 34– 48 for all time points). Kruskal-Wallis test with Dunn’s multiple comparisons. Columns labeled with different letters are significantly different from each other (P < 0.05). (**C**) Total activity 6–10 min post-CO2 under LD entrainment in the photophase (n = 34–48 for all time points). Kruskal-Wallis test with Dunn’s multiple comparisons. Columns labeled with different letters are significantly different from each other (P < 0.05). (**D**) Average duration of activity bouts under LD entrainment in the photophase (n = 34–48 for all time points). Ordinary one-way ANOVA with Tukey’s multiple comparisons test. Columns are not significantly different from each other (P > 0.05). (**E**) Total initial CO2-evoked activity (0–2 min post-CO2) on the second DD cycle (n = 20 for all time points). Kruskal-Wallis test with Dunn’s multiple comparisons. Columns are not significantly different from each other (P > 0.05). (**F**) Total sustained CO2-evoked activity (2–10 min post-CO2) on the second DD cycle (n = 20 for all time points). Kruskal-Wallis test with Dunn’s multiple comparisons. Columns are not significantly different from each other (P > 0.05). (**G**) Average duration of activity bouts on the second DD cycle (n = 20 for all time points). Ordinary one-way ANOVA with Tukey’s multiple comparisons test. Columns are not significantly different from each other (P > 0.05). Shaded lines show mean ± SEM. White and dark grey shading in **A–D** represent light and dark phases, respectively. Light and dark grey shading in **E–G** represent respective light and dark phases from prior LD entrainment, respectively.

**Data S1. (separate file)**

All raw data and statistical test details used for main and supplemental figures, with the exception of fig. S6 and S7, which is included in Data S2. This file also include plasmid sequence used for the generation of the *pdf-QF2* mutant (see Material and Methods).

**Data S2. (separate file)**

Behavioral classification of animals in the MozzieDrome circadian host-seeking assay, raw data used for fig. S6 and S7.

